# Tibicos differentially affects faecal microbiota composition and short-chain fatty acid production in prediabetic and healthy adults in a simulated digestive tract

**DOI:** 10.1101/2025.03.26.645606

**Authors:** Miin Chan, Xinwei Ruan, Jing Wang, Liz Leya Ninu, Kate Howell

## Abstract

Prediabetes and obesity are increasingly prevalent chronic pro-inflammatory conditions driven by gut microbiota dysfunction. To assist in the management of such diseases, researchers are interested in gut-microbiota targeted dietary interventions. Tibicos is a plant-based fermented beverage shown to modulate gut microbiota, with anti-hyperglycaemic and anti-inflammatory effects in *in vitro* and *in vivo* human and animal studies. We investigated the effects of ginger-cayenne tibicos on healthy and prediabetic pooled faecal microbiota, with a static *in vitro* digestion/ colonic fermentation model using faeces from adult overweight/ obese donors with prediabetes, and healthy donors (*n*=3 per group). Changes in faecal microbiota composition and short-chain fatty acid (SCFA) production were determined. Baseline prediabetic faeces was less abundant in butyrate-producing bacterial taxa and SCFAs. *In vitro* colonic fermentation Time and Status (prediabetic vs. healthy) were the major determinants of shifts in faecal microbial community abundance and diversity. Tibicos reduced beneficial bacterial taxa loss in prediabetic faecal microbiota. However, tibicos improved the functional capacity of healthy faecal microbiota to a larger extent, with increased SCFA production and significant correlations between bacterial species and SCFAs. The differential shifts in composition and SCFAs observed during simulated gastrointestinal transit suggest that ginger-cayenne tibicos has the potential to beneficially modulate human faecal microbiota in both healthy and prediabetic participants. This study indicates that ginger-cayenne tibicos may be a useful tool in the management of prediabetes and obesity in humans and provides pilot data for the development of larger clinical trials.

## Introduction

The human gut microbiota plays a central regulatory role in immune function and energy metabolism ^(1)^. Diet has a major impact on human health, mainly through the interaction between food components and the gut microbiota ^(2,3)^. Poor diets are associated with dysfunctional gut microbiota, and both are linked to poorer health outcomes ^(4)^. These complex relationships underlie the development of chronic metabolic inflammatory diseases such as type 2 diabetes mellitus (T2DM), metabolic syndrome (MetS) and cardiovascular disease. Two highly prevalent components of MetS, prediabetes and obesity, are significant cardiovascular risk factors, predisposing individuals to developing T2DM: up to 10% of people per year with prediabetes will develop diabetes ^(5)^The global prevalence of prediabetes in 2021 was 5.8% (298 million), with a projected increase to 6.5% (414 million) by 2045 ^(6)^; 1 billion adults will develop obesity by 2030, with T2DM projected to affect 700 million adults by 2045 ^(7)^. As such, researchers are developing gut-microbiota-targeted dietary strategies to manage prediabetes and obesity.

Botanical fermented foods (BFFs) are produced through the action of bacteria and yeast on vegetables, fruit, cereals, nuts and pulses ^(8)^. BFFs are known to be rich in health-promoting components, including food-associated microorganisms and their metabolic products, dietary fibre, antioxidants and phenolic compounds ^(9)^. Compared to their dairy-based counterparts, these foods contain more diverse microbial communities, with more microbial genes associated with potential health benefits to the host ^(10)^. A randomised prospective clinical trial showed that high levels of BFF consumption increased microbiota diversity and reduced inflammatory markers, when compared to a high-fibre diet ^(11)^. Cheap to produce and already globally consumed, BFFs are increasingly studied for the beneficial manipulation of the microbiota-gut-metabolism axis ^(1)^. A myriad of BFFs remain under-studied, with researchers focused on uncovering their functional effects for the prevention and management of disease.

Tibicos, also known as water kefir, is a traditional Mexican low-alcohol fermented beverage. Despite the purported health benefits of tibicos, there remains a dearth of evidence regarding its effects on the gut microbiota and human health. Tibicos is well-established as a source of dietary microorganisms, including lactic acid bacteria (LAB), acetic acid bacteria, and yeasts ^(12–14)^. Its functional abilities are likely due to the antimicrobial activity of its microbially-produced bioactive compounds ^(15)^, via the direct (transient gut integration) and indirect (short-chain fatty acid [SCFA]/ metabolite production) actions of LAB and other microorganisms found in the tibicos liquor ^(16,17)^. Recent metagenomic studies have further identified bacterial and yeast species in tibicos, along with functional analysis linking these microbes to the production of organic acids ^(13)^. Leech et al. ^(10)^ found that tibicos was significantly more microbially diverse than dairy-based ferments, with the largest number of potential health-promoting gene clusters. *In vitro* tests of yeast and lactobacilli strains isolated from tibicos showed that they highly adhered to intestinal cells; had good resistance to low gastric pH, bile salts and *in vitro* digestive fluids; and possessed antioxidant properties ^(18,19)^. *In vivo* studies in artificially diseased Wistar rats have indicated tibicos’ hepatoprotective, anti-inflammatory, antiulcerogenic and DNA protective/ antioxidant properties ^(20)^. Rats with streptozotocin-induced diabetes supplemented with tibicos lost weight and had improved plasma glucose and lipid profiles compared to control, with antihypertensive effects ^(21,22)^. Notably, there are no human clinical trials with tibicos as an intervention in healthy humans or in our population of interest ^(23)^. Before advancing to clinical trials, more appropriate pilot data is required, which can be collected *in vitro* ^(24)^ without the need for expensive *in vivo* studies.

In our previous *in vitro* digestion study ^(8)^, we administered ginger and cayenne tibicos to healthy porcine faeces. We confirmed other research which showed that tibicos contains probiotic levels (6 to 7 log CFU/g) of viable microorganisms ^(25)^, and observed that microbes from tibicos had high survival rates after gastrointestinal (GI) transit, significantly improved by the addition of ginger and cayenne pepper (*p*<0.05). We also found that tibicos shifts faecal microbiota composition, with an increased relative abundance of potentially beneficial taxa such as *Megasphaera*. Calatayud et al.’s ^(17)^ *in vitro* digestion study using healthy human faecal donors found that tibicos increased beneficial SCFA production by microbes, beneficial *Bifidobacterium* abundance, and anti-inflammatory cytokines that strengthened the intestinal barrier. Notably, these two studies are the only ones that have tested tibicos (as opposed to microbes isolated from tibicos) with human faeces. The aim of this study was to investigate the impact of ginger cayenne tibicos on faecal microbiota composition (abundance and diversity) and function (SCFA production) in overweight/ obese adults with prediabetes, compared to healthy controls. To achieve this, we used a validated *in vitro* digestion and colonic fermentation model ^(26,27)^ combined with 16S rRNA long-read sequencing, quantitative PCR, bioinformatics, and GC-FID.

## Methods

### Chemicals and reagents

Fresh tibicos grains were donated by Dr. Miin Chan. Organic dried figs were purchased from Woolworths (Melbourne, VIC, Australia). Organic ginger powder and cayenne pepper were purchased from Queen Victoria Market (Melbourne, VIC, Australia). DNA was extracted using DNeasy PowerSoil Pro kits from Qiagen (Clayton, VIC, Australia). The following analytical grade reagents and chemicals were purchased from ChemSupply (Gillman, SA, Australia): sucrose, propionic acid, orthophosphoric acid, calcium chloride, sodium hydroxide, sodium acetate, magnesium chloride hexahydrate, ammonium chloride, potassium chloride, sodium phosphate dibasic, and potassium sodium tartrate tetrahydrate. From ThermoFisher (Scoresby, VIC, Australia), glacial acetic acid and formic acid were purchased. Analytical grade reagents and chemicals were purchased from Merck (Bayswater, VIC, Australia): tris(hydroxymethyl)aminomethane, peptone, potato starch, ⍺-amylase from human saliva, haemoglobin from bovine blood, pepsin from porcine gastric mucosa, trichloroacetic acid, N⍺-p-Tosyl-L-arginine methyl ester hydrochloride, sodium bicarbonate, cycloheximide, 3,5-Dinitrosalicylic acid, resazurin sodium salt, L-Cysteine, sodium sulfide hydrate, 4-methyl valeric acid, and butyric acid. Pancreatin was purchased from MP Biomedicals (Seven Hills, NSW, Australia), whilst Carrez clarification and bile acid assay kits were purchased from Abcam (Melbourne, VIC, Australia).

### Ginger cayenne tibicos preparation, microbial enumeration and pH

Tibicos was aseptically prepared in Schott bottles with airlocks, as described in Chan et al. ^(8)^. Briefly, after primary fermentation at room temperature (20-25 °C) for 72h, the liquor was transferred to another sterile 1L Schott bottle. 0.5% (w/v) organic ginger (*Zingiber officinale*) and 0.125% (w/v) organic cayenne (*Capsicum frutescens*) powder were added to the liquor for secondary fermentation. After flushing with nitrogen, the ginger cayenne tibicos was incubated at room temperature for 48h then stored at 4 °C until *in vitro* digestion. Samples were taken on day 0, 1, 3, 5 and 7 for LAB enumeration, which was performed using the methods of Laureys and De Vuyst ^(28)^ with modifications. Serial dilutions (10^-1^ to 10^-6^) of 1 mL liquor samples in 0.9 % (w/v) sterile peptone solution were prepared and plated on MRS agar supplemented with 4 mg/L cycloheximide. Plates were anaerobically incubated at 27 °C for 3 days before colony enumeration. pH was determined using an Edge pH Benchtop kit (Keysborough, VIC, Australia).

### Prediabetic and healthy control faeces collection and standardisation

Faecal samples from healthy individuals were provided by four female participants of East Asian descent, aged between 18 to 25, with an average body mass index (BMI) of 21.9 kg/m^2^. Faecal samples from adults with prediabetes, as per the American Diabetes Association ^(29)^, were provided by two female participants of Anglo-Saxon descent, aged between 30 to 45, with an average HbA1c of 6.1%, and BMI of 33.5 kg/m^2^. The participants were all nonsmokers in good health who had not consumed antibiotics, probiotics, prebiotics, laxatives, metformin, glucagon-like peptide-1 agonists, or other antidiabetic/ weight loss medications in the three months prior to collection. Participants were not regular consumers of fermented foods, but otherwise consumed an unrestricted diet. Fresh faecal samples were collected in airtight sterile specimen containers, kept below 4 °C and received within 12 hours. Upon receipt, in a biosafety cabinet, samples were vortexed 50:50 with 20% (wt/vol) glycerol, snap frozen with liquid nitrogen and stored at –80 °C until commencement of *in vitro* colonic fermentation as per Pérez-Burillo et al. ^(27)^.

### Human ethics, data storage and privacy

This study was conducted according to the guidelines laid down in the Declaration of Helsinki (1975, revised in 2008) and all procedures involving human subjects were approved by the University of Melbourne Human Research Ethics Committee (Ethics approval number 2024-27258-55593-7.). Participation was voluntary and after a briefing session, all participants signed an electronic informed consent form and questionnaire using Qualtrics Software, Version 2024 (Qualtrics, Provo, Utah, USA). A random number generator was used to assign a random four-digit number to each participant. Identifying information was not used in publications or databases.

### Simulated upper GI digestion and colonic fermentation

To simulate the digestion of tibicos in the human upper gastrointestinal tract, *in vitro* digestion was performed as per the INFOGEST protocol ^(26)^ and Minekus et al. ^(30)^ with minor modifications, as described below. *In vitro* colonic fermentation followed the procedure described in Pérez-Burillo et al. ^(27)^. Tibicos samples were taken on day 21, with the microbial load of approximately 7 log CFU/mL, as per our previous study ^(8)^, for use in the *in vitro* digestive system. This time point was chosen to maximise the microbial load at time of administration, as well as consider storage length in a commercial context ^(8)^. All experiments were conducted in triplicate.

## Simulated *in vitro* digestion

Simulated salivary fluid (SSF), simulated gastric fluid (SGF), simulated intestinal fluid (SIF), and digestive enzymes (α-amylase, pancreatin, pepsin and bile) were prepared according to methods described in Brodkorb et al. ^(26)^, including enzyme activity assays. These were used fresh, i.e. within an hour of preparation. Blanks using Milli-Q water instead of tibicos were also produced.

Oral digestion: 5 mL of tibicos (or Milli-Q water for the blanks) was mixed with 3.5 mL of SSF solution, 0.5 mL human salivary α-amylase solution (75 U/mL), 25 µL of 0.3 M CaCl_2_(H_2_O)_2_ and 925 µL of Milli-Q water. The final sample-to-stock ratio was 1:1. Tubes were tightly sealed and placed in a prewarmed 37 °C shaking incubator for 2 min at 200 rpm to obtain an oral bolus.

Gastric digestion: 10 mL of the oral bolus was added to 7.5 mL of SGF solution, 1.6 mL of porcine pepsin solution (25000 U/mL), 5 µL of 0.3 M CaCl_2_(H_2_O)_2_ and 695 µL of Milli-Q water. The final sample-to-stock ratio was 1:1. pH was adjusted to 3.0 with required amount of 1 mol/L HCl. Sealed tubes were placed into a prewarmed 37 °C shaking incubator for 2h at 150 rpm to obtain gastric chyme.

Small intestinal digestion: 20 mL of gastric chyme was mixed with 11 mL of SIF solution, 5 mL pancreatin solution (800 U/mL), 2.5 mL bile solution (49 mg/mL), 40 µL of 0.3 M CaCl_2_(H_2_O)_2_ and 1.31 mL of Milli-Q water. pH was adjusted to 7.0 with required amount of 1 M NaOH. Sealed tubes were placed into a prewarmed 37 °C shaking incubator for 2h at 150 rpm. Resulting samples were stored at –20 °C for analysis and *in vitro* colonic fermentation.

## *In vitro* colonic fermentation

Phosphate buffer, fermentation medium and faecal slurry were freshly prepared in aseptic conditions according to methods described in Pérez-Burillo et al. ^(27)^. Controls contained the contents of the blank tube from *in vitro* digestion, fermentation medium and faecal slurry (no tibicos).

The fermentation medium was prepared as per Pérez-Burillo et al. ^(27)^. Briefly, in a biosafety cabinet, 1 L of pH-adjusted (7.0) peptone solution was mixed with 50 mL of reductive solution (containing cysteine and sodium sulfide) and 1.25 mL of 0.1% (w/v) resazurin solution for each litre of fermentation medium.

For faecal slurry preparation, faecal samples (stored at –80 °C) were thawed in a 37 °C water bath for 30 min. Glycerol was removed by centrifugation at 4000 g for 10 min at 4 °C. Supernatant was discarded, and the pellet retained. 53.8 g (13.5 g per participant) of the faecal pellet was used for the healthy group, and 40.32 g (10.08 g per participant) for the prediabetic group. Faecal pellets were transferred to stomacher bags and resuspended in autoclaved room temperature 0.1 M phosphate buffer at a faecal concentration of 32% (w/v), then homogenised with a vortex for 1 min. Homogenised faecal slurry was distributed into three tubes per group and centrifuged at 550 g for 5 min at room temperature. The resulting supernatant was used for *in vitro* colonic fermentation.

To commence *in vitro* colonic fermentation, all solutions were prewarmed to 37 °C. 1 mL of small intestinal digesta (samples or blanks) was added to 7.5 mL of fermentation medium and 2 mL of faecal slurry. All tubes were flushed for 10 seconds with nitrogen, sealed tightly, vortexed and placed in a prewarmed 37 °C shaking incubator for 24h at 100 rpm. 1 mL aliquots were collected from different tubes at five time points (0, 4, 8, 12, and 24h) during colonic fermentation, snap frozen with liquid nitrogen and stored at –20 °C for further analysis.

### Sample and microbial DNA extraction

Bacterial genomic DNA was extracted in triplicate from 200 μl of sample using the DNeasy® PowerSoil® Pro Kits (QIAgen, CA, USA), as per the manufacturer’s instructions. The quality of extracted DNA was verified by electrophoresis on a 1.2% (w/v) agarose gel. Controls (faecal slurry with digestive fluids and Milli-Q water replacing tibicos before and after *in vitro* fermentation) were included. Tibicos, small intestinal digesta and faecal samples were directly used for extraction. Precipitates from *in vitro* colonic fermentation samples were thrice washed with ice-cold phosphate buffer saline solution (with 2% v/v polyvinylpolypyrrolidone) at a ratio of 2:3, removing any phenolic material. The subsequent pellets were mixed with DNA/ RNA shield solution before performing DNA extraction. The concentration and quality of extracted genomic DNA were evaluated using the NanoDrop ND2000c spectrophotometer (NanoDrop Technologies, Wilmington, DE, USA) and standardised to 1 ng/μL prior to performance of PCR.

### Microbial quantification with quantitative PCR

Quantitative PCR analysis was performed to determine the total gene copy number (GCN) per µL of each sample, using the universal bacterial primer set Eub 338/Eub518. DNA extracts were sent to TrACEES (The University of Melbourne, Parkville, Australia) for qPCR analysis, as per their standard protocol ^(31,32)^. Briefly, the 36 extracted DNA samples, including controls and triplicates, underwent an “all bacteria” assay. This was achieved using a BioRad MyiQ Single-Color Real-Time PCR Detection System in 25 μl reactions containing 12.5 μl iQ SYBR Green Supermix (BioRad, South Granville, NSW, Australia); 2 μl template DNA (1:1000 diluted), 300 nM of both primers 16S-341_F and 16S-805_R. Negative (DNA replaced by sterilised water in qPCR) and positive (quantified recombinant plasmid with the target 16S rRNA sequence) controls were included in each run, as well as the controls for each group (faecal slurry with digestive fluids and Milli-Q water). The PCR amplification program consisted of an initial denaturation set at 95℃ for 3 min, followed by 40 cycles at 95℃ for 5 sec and 60℃ for 30 sec. Estimated absolute abundance of 16S rRNA genes across different groups and time points was determined by comparison with automatically created standard curves (Bio-Rad software) generated by preparing 10-fold serial dilutions of plasmids.

### Microbial community analysis with long-read 16S rRNA amplicon sequencing

DNA extracts were submitted to the Australian Genome Research Facility for PacBio HiFi sequencing as per the PacBio protocol. Briefly, the concentration of genomic DNA was quantified with the Qubit HS DNA kit (Thermo Scientific™) and normalised to 1 ng/ μL, Full-length 16S amplicons were generated using Platinum™ SuperFi II High-fidelity PCR enzyme (Thermo Scientific™) from 2 ng of genomic DNA. Amplicons were pooled and purified with single-molecule real-time (SMRT)Bell™ cleanup beads (PacBio). The resulting amplicon library was contracted using the SMRTBell™ 3.0 library prep kit (PacBio), polymerase-bounded with the Revio Polymerase Kit (PacBio) and loaded into a SMRT Cell 25M. 16S rRNA full-length sequencing (V1 to V9) was performed on the PacBio Revio instrument for 30 hours.

### SCFA analysis with gas chromatography-flame ionisation detection

The analysis of short-chain fatty acids (SCFAs) acetate, butyrate and propionate was conducted as described by Gu et al. ^(33)^. Gas chromatography (GC) was coupled with a flame ionisation detector (FID), an autosampler, and an autoinjector (Agilent, CA, USA). A SGE BP21 capillary column and a retention gap kit (SGE International, Ringwood, VIC, Australia) were used. Helium served as the carrier gas, with a flow rate of 14.4 mL/min. GC conditions were as follows: 100°C for 30 s; 180°C for 1 min (6°C/min); 200°C for 10 min (20°C/min). FID temperature and injection port temperature were set at 240°C and 200°C, respectively. Makeup gases included nitrogen, hydrogen, and air, with flow rates of 20, 30, and 300 mL/min, respectively. Sample injection volume was 1 µL. All results were converted to mmol/L per digesta for subsequent statistical analysis.

### Statistical analysis and bioinformatics

Statistical analyses were performed using GraphPad Prism v10.3.1 (GraphPad Software, Boston, MA, USA). Differences in total GCN and SCFA concentration among samples, at various fermentation timepoints where relevant, were analysed with one-way analysis of variance (ANOVA), with repeated measures where appropriate, followed by Tukey’s HSD test. Significance level was set at *p*<0.05. All results are expressed as the mean ± standard deviation. Collected faecal samples were homogenised into two groups, then sampled and tested in triplicate; all other experiments were performed in triplicate.

For microbiome bioinformatic analyses, demultiplexed PacBio HiFi full-length 16S rRNA gene sequences were imported and processed using QIIME2 ^(34)^. Quality filtration and deionisation into amplicon single variants (ASV) were performed with DADA2 ^(35)^, using the function “dada2 denoise-ccs”. Taxonomic classification was achieved via consensus alignment classification with VSEARCH against the Genome Taxonomy Database (GTDB r207) ^(36)^, as well as naïve Bayesian machine learning based classification i.e. DADA2. If a species-level match was not identified, further classification was achieved with three databases that successively referred to the next one: GTDB r207, SILVA v138 ^(37)^ and the NCBI RefSeq 16S rRNA database supplemented by the Ribosomal Database Project. The phylogenetic tree was constructed by the *mafft* alignment. Data exported from QIIME2 were further analysed in R (version 4.2.2).

For diversity analyses, sequencing data was rarefied to 10,000. α-diversity of bacterial communities was calculated using Shannon’s diversity index and Chao1. Statistically significant (*p*<0.05) differences between groups were determined using the factorial Kruskal−Wallis sum rank test (α=0.05), with post-hoc analysis performed with Dunn’s test. For β-diversity, unweighted UniFrac distances between samples were calculated and demonstrated with principal coordinate analysis (PCoA). Adonis2 function of vegan package ^(38)^ was used to perform permutational multivariate analysis of variance (PERMANOVA) with 999 permutations, which determines statistically significant (*p*<0.05) differences in beta-diversity. Spearman’s correlations were calculated to assess the relationship between the relative abundance of bacterial taxa (mean relative abundance >0.5%) and SCFAs and visualised using the *corrplot* package ^(39)^. Analysis of microbiome composition with bias correction 2 (ANCOM-BC2) ^(40)^ was performed on count numbers at genus and species level to identify differential taxa. Taxa showing differential abundance between groups were visualised with the *ComplexHeatmap* R package ^(41)^.

## Results

Results are ordered to represent tibicos moving through the upper and lower GI tract. Microbial composition (relative abundance and total gene copy number) of tibicos, faecal samples and colonic fermenta are presented first, followed by function (SCFAs). Lastly, these results are differentially analysed to produce β– and α-diversity, along with a combined clustered/ ANCOMBC-2 heatmap and bacterial species/ SCFA correlation plot.

### Microbial composition and quantification of ginger-cayenne tibicos and healthy/ prediabetic faeces

#### *In vitro* digestion significantly decreased bacterial content in tibicos, whilst *in vitro* fermentation increased bacterial content in faecal microbiota

The total gene copy number (GCN) of tibicos decreased significantly (*p*=0.001) after *in vitro* digestion (Supplementary material 1, Figure 1A). There were no significant differences between the total GCN between healthy (H) and prediabetic (P) faecal samples (Supplementary material 1, Figure 1B). 24h of *in vitro* fermentation led to an increase in total GCN in all samples (Supplementary material 1, Figure 1C). In the healthy fermenta, there was a significant increase (*p*=0.0294) in total GCN from 0h to 24h, as well as in the healthy control (*p*=0.0095). In both the prediabetic fermenta and prediabetic controls, the increase in total GCN was not significantly different between the two timepoints. There were no significant differences between each group and their control groups at each time point. When comparing the healthy and prediabetic fermenta, there was no significant difference at any time point.

**Figure 1.**
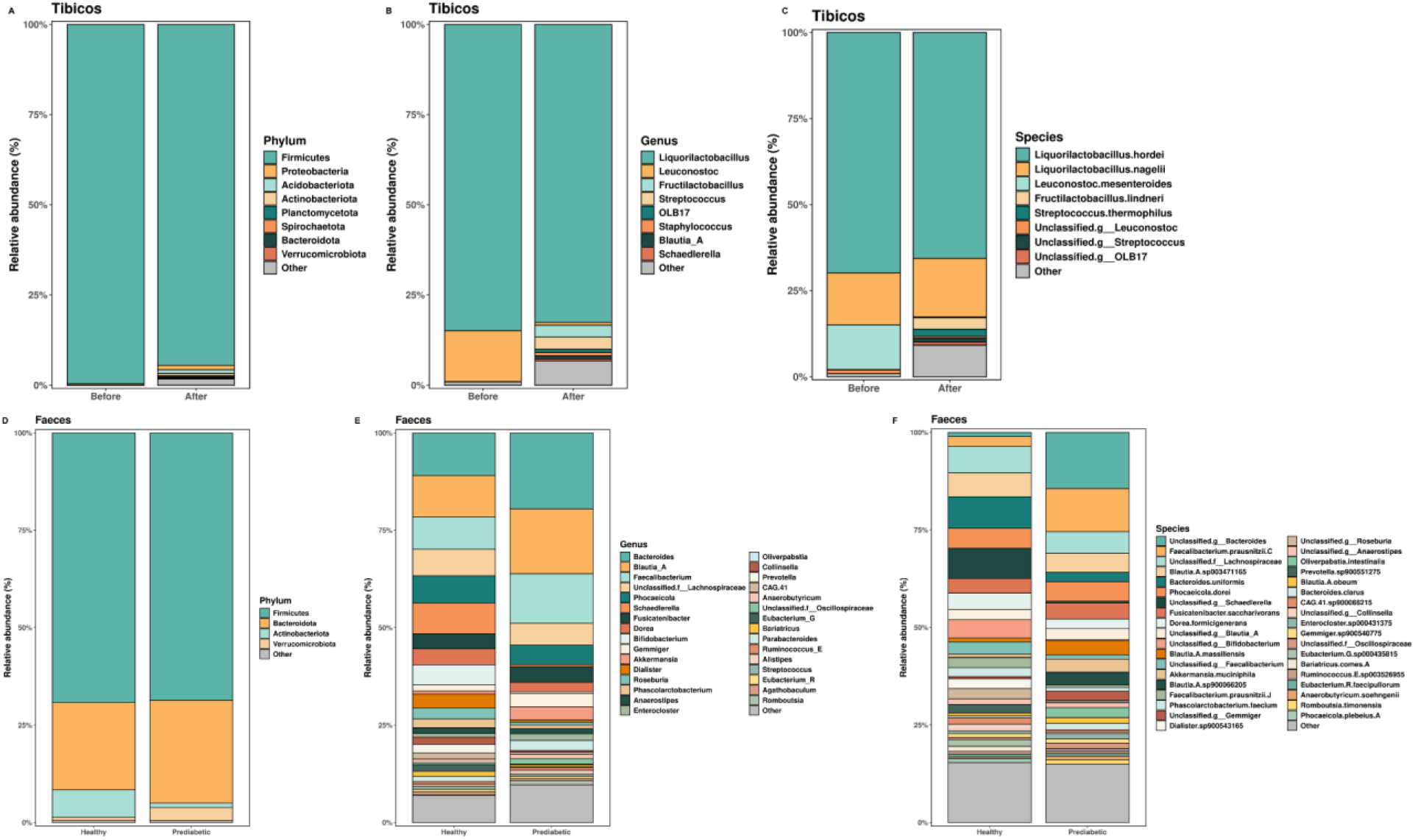
Ginger-cayenne tibicos microbial composition is affected by *in vitro* digestion, and the microbial composition of healthy and prediabetic faeces differs in relative abundance. Microbial taxa in **ginger-cayenne tibicos** before and after *in vitro* digestion are classified by **(A)** phyla **(B)** genus, and **(C)** species levels (highest 8 MRA). Before: ginger-cayenne tibicos only; After: ginger-cayenne tibicos and digestive enzymes post-*in vitro* digestion. Microbial taxa in **healthy and prediabetic faecal samples** are classified by **(D)** phyla, **(E)** genus, and **(F)** species levels (mean relative abundance > 0.5%); All results are ordered according to abundance and expressed as mean ± SD (*n*=3).

#### *In vitro* digestion causes minor shifts in ginger-cayenne tibicos microbial composition

In ginger-cayenne tibicos, *Firmicutes* was the dominant phyla (Figure 1A), with a small drop observed after *in vitro* digestion. Some phyla were only observed post-digestion, including *Acidobacteriota* and *Actinobacteriota*. At the genus level (Figure 1B), *Liquorilactobacillus* dominated, with a small drop post-digestion; *Leuconostoc* declined by a factor of 18. Other genera were only observed after *in vitro* digestion, including *Fructilactobacillus*, *Streptococcus*, *OLB 17*, *Staphylococcus*, *Blautia A* and *Schaedlerella*. There were observed reductions in two dominant species (Figure 1C), *Liquorilactobacillus hordei* and *Leuconostoc mesenteroides*. Otherwise, most changes in abundance were due to increases observed post-digestion: *Liquorilactobacillus nagelii*, *Fructilactobacillus lindneri*, and *Streptococcus thermophilus*. *Unclassified g_Leuconostoc* species, *Unclassified g_OLB17, Akkermansia muciniphila* and *Faecalibacterium prausnitzii* only appeared post-digestion. Mean relative abundance at class, family and order levels are available in the Supplementary material (Supplementary material 1, Figure 2 and Supplementary material 2).

**Figure 2.**
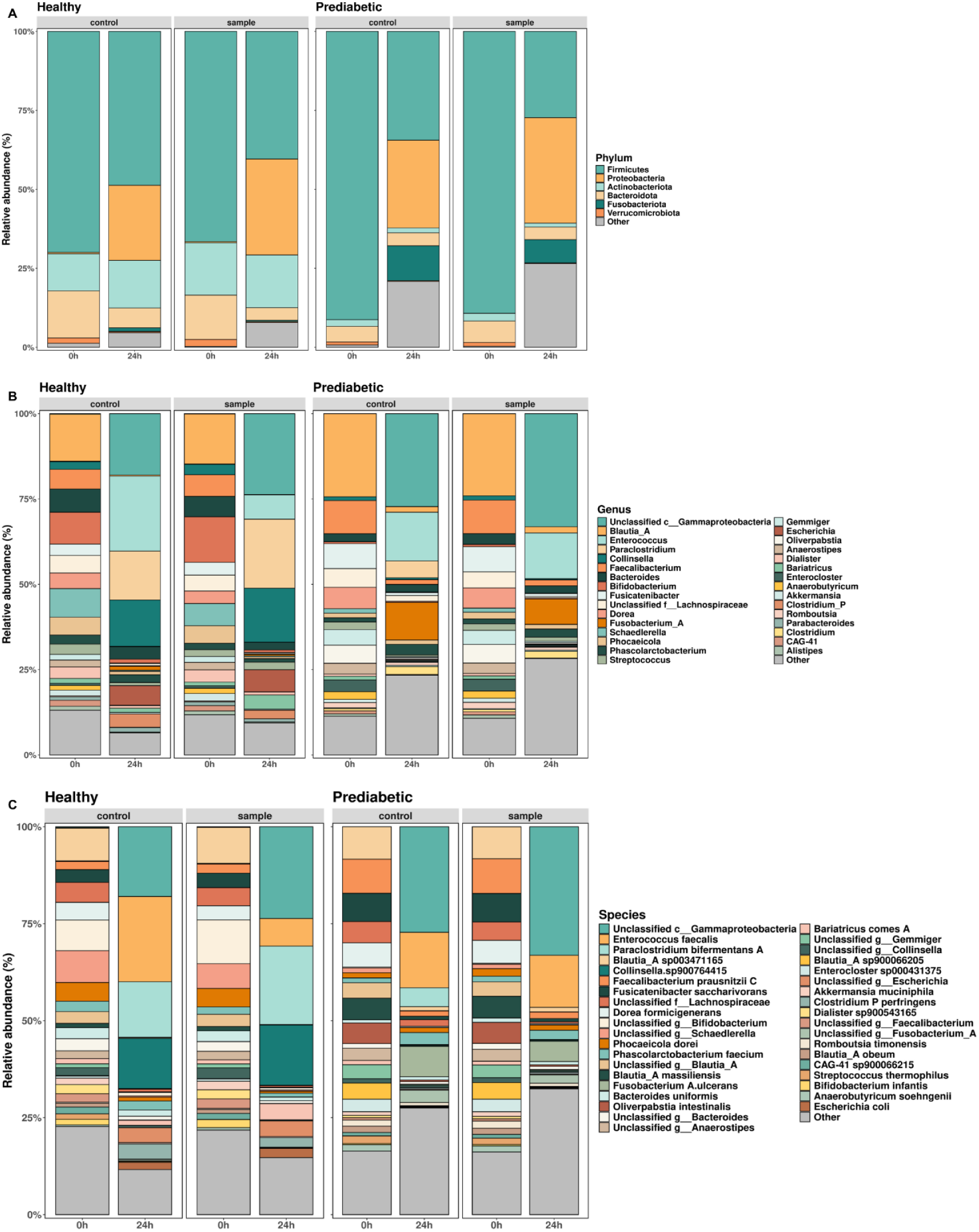
*In vitro* colonic fermentation changes faecal microbiota composition in healthy and prediabetic faeces. Relative abundance plots were generated using the most abundant bacterial taxa (>0.5%). Microbial taxa are classified by (A) phyla, (B) genus, and (C) species levels. Healthy and prediabetic samples: fermentation media + faecal slurry + tibicos digesta; healthy and prediabetic controls: fermentation media, faecal slurry and control digesta. Results expressed in mean ± SD (*n*=3).

#### Healthy and prediabetic faeces have distinct microbial compositions

In both healthy and prediabetic faeces, *Firmicutes* was the dominant phyla (Figure 1D). *Bacteroidetes* was similarly abundant in both groups as was *Proteobacteria*. Compared to prediabetic faeces, healthy faeces was more abundant in *Actinobacteriota*, but less so in *Verrucomicrobiota*. Accordingly at the genus level (Figure 1E), *Bacteroides* and *Blautia A* were most abundant in both faecal samples but were approximately twice as abundant in prediabetic faeces. Prediabetic faeces was also more abundant in *Faecalibacterium*, *Akkermansia*, *Gemmiger*, *Oliverpabstia* and others. However, healthy faeces was more abundant in many key genera involved in beneficial SCFA production, including *Schaedlerella*, *Phoecaeicola*, *Bifidobacterium*, *Dorea*, *Dialister*, *Roseburia*, *Prevotella* and *Phascolarctobacterium*. *Streptococcus, Unclassified.f_Lachnospiraceae* and *Fusicatenibacter* were similarly abundant in both groups. At the species level (Figure 1F), *Unclassified.g_Bacteroides* was the most dominant species in prediabetic faeces and was much more abundant than in healthy faeces. Prediabetic faeces was five times more abundant in *Faecalibacterium prausnitzii C*, as was *Unclassified.g_Gemmiger*, whilst *A. muciniphila* was four times more abundant. Healthy faeces was more abundant in *Bacteroides uniformis*, *Unclassified.g_Bifidobacterium*, *Unclassified.g_Schaedlerella*, *Prevotella sp900551275*, *Unclassified.g_Roseburia*, *Dialister sp900543165*, *Dorea formicigenerans*, *Phasoclarctobacterium faecium* and *Roseburia intestinalis*. Both faecal samples had similar abundance of Unclassified.f_*Lachnospiraceae*, *B. A sp003471165*, *Phocaeicola dorei* and *Fusicatenibacter saccharivorans*. Mean relative abundances at class, family and order levels for healthy and prediabetic faeces are available in the Supplementary material (Supplementary material 1, Figure 3 and Supplementary material 2).

**Figure 3.**
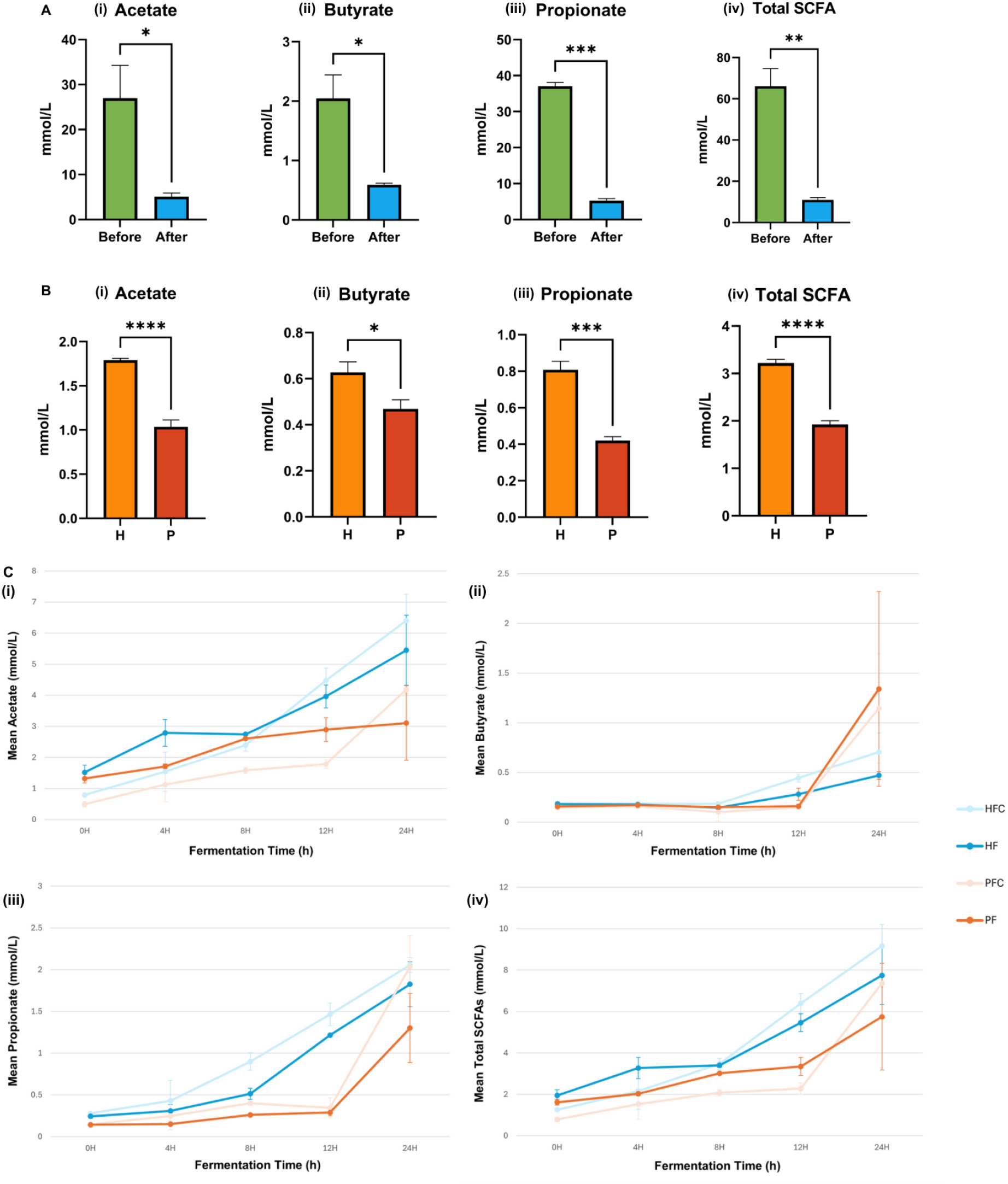
*In vitro* digestion of tibicos reduces short-chain fatty acid concentrations, whilst *in vitro* colonic fermentation increases concentrations. **(A) Changes in tibicos sample before and after *in vitro* digestion:** (i) Acetate; (ii) Butyrate; (iii) Propionate; and (iv) total short-chain fatty acids. Results were compared using paired t-tests. **(B) Differences between healthy and prediabetic faecal samples:** H: healthy faecal samples; P: Prediabetic faecal samples. Results were compared using unpaired t-tests. Significant differences between groups * *p*<0.05 ** *p*<0.01 *** *p*<0.001 **** *p*<0.0001. **(C) Changes in healthy and prediabetic fermenta during 24h *in vitro* colonic fermentation:** HF: fermenta containing healthy faecal slurry; PF: fermenta containing prediabetic/ obese faecal slurry; HFC: healthy fermentation control; PFC: prediabetic fermentation control. Control tubes contained *in vitro* digestion control (MilliQ water replacing tibicos) mixed with respective faecal slurries. All results expressed as mean ± standard deviation (*n* = 3).

### Changes in microbial composition of healthy and prediabetic fermenta during *in vitro* colonic fermentation with ginger-cayenne tibicos

#### Faecal microbiota composition of healthy and prediabetic fermenta were different to each other, and were changed by *in vitro* colonic fermentation with ginger-cayenne tibicos

To compare the faecal microbial communities of healthy and prediabetic participant groups, the full-length 16SrRNA gene was sequenced. 2611 amplicon sequence variants (ASV) were identified. As per our earlier findings, the healthy and prediabetic fermenta had differing microbial compositions, with subtle differences to their own controls (containing Milli-Q water instead of tibicos) at 0h, and larger differences at 24h (Figures 2A to C). *In vitro* colonic fermentation led to similar patterns of change in all groups between the start (0h) and end of colonic fermentation (24h).

In comparison to their controls, both groups had small differences in relative abundance at 0h, especially noticeable at genus/ species levels (Figures 2B, C). In the healthy fermenta, at 0h, *Bifidobacterium* was more abundant in the sample vs. the control; by 24h, *Bifidobacterium* was reduced in both, but it was 1.5 times less abundant in the sample vs. the control. *Enterococcus faecalis* was the same in the healthy control and sample at 0h, but by 24h was three times more abundant in the control. The opposite was observed with *Unclassified.c_Gammaproteobacteria*, and *Paraclostridium bifermentans A*, with both becoming more abundant in the samples compared to controls. In the prediabetic fermenta, there were less differences observed between controls and samples. *Unclassified.c_Gammaproteobacteria* was similar in the control and sample at 0h, but at 24h was more abundant in the sample. *P. bifermentans A* was not detected at 0h in either prediabetic control or sample, but at 24h was at five times higher in the control, whilst remaining undetectable in the sample. Although these differences are subtle, they do indicate that ginger-cayenne tibicos has an effect on microbial composition compared to control, especially after colonic transit.

As established above, there were many differences in microbial composition on all taxonomy levels between the faecal samples. However, when these samples were used to produce the fermenta groups, differences in microbial composition between the groups appeared, likely due the chemical process involved. For example, *Firmicutes* was more abundant in the prediabetic fermenta, and *Bacteroidetes* in the healthy fermenta, even though they were the same abundance in the pure faecal samples. The patterns of change over 24h differed between the two fermenta groups, with upward or downward shifts in mean relative abundance, some by larger factors than others. In both fermenta groups, after *in vitro* fermentation with ginger-cayenne tibicos, there was a decline in abundance of *Firmicutes* (healthy 1.6x; prediabetic 3.2x), *Bacteroidetes* (healthy 3.5x; prediabetic 1.7x) and *Verrucomicrobiota* (healthy 8x; prediabetic 6x) (Figure 2A). However, *Actinobacteria* abundance remained the same in healthy fermenta, whilst dropping in the prediabetic fermenta. Large increases in abundance were observed in *Proteobacteria* (healthy 82 x; prediabetic 740x), and *Unclassified.d_Bacteria* (healthy 90 x; prediabetic 82 x). In both groups. *Fusobacteriota* only made an appearance post-fermentation, and much more so in healthy fermenta.

Certain genera were reduced by *in vitro* colonic fermentation (Figure 2B). These include *Blautia A* (healthy 98x; prediabetic 13x); *Bifidobacterium* (healthy 20x; prediabetic 5x); *Dorea* (healthy 22x; prediabetic 18x); and *Fusicantenibacter* (healthy 41x; prediabetic 9x). Also reduced at 24h were *Dialister* and *Roseburia*, which were more abundant in the healthy fermenta, and *Gemmiger,* higher in the prediabetic fermenta. *Akkermansia* was similar in both fermenta and was reduced by fermentation, as were *Unclassified.f_Lachnospiraceae*, *Faecalibacterium*, *Schaedlerella*, *Bacteroides* and *Phocaeicola*. Other genera increased in abundance by 24h. *Unclassified.c_Gammaproteobacteria* was barely detectable at 0h but became the dominant genera in both fermenta (healthy 236x; prediabetic 1657x) at 24h. *Collinsella* in healthy fermenta was increased post-fermentation and reduced in the prediabetic fermenta. *Unclassified.d_Bacteria* and *Enterococcus* increased in both groups post-digestion, but more so in the prediabetic fermenta. At 24h, no *Paraclostridium* was detected in prediabetic fermenta, whilst in the healthy fermenta it increased by a factor of 20 Similarly, *Escherichia* remained at around 0 in the prediabetic fermenta, but increased 214 times in healthy fermenta. Notably, there was a lack of *Lactobacillus*, the dominant genera in tibicos, across timepoints and groups.

Species reduced by 24h of *in vitro* colonic fermentation (Figure 2C) in both fermenta groups include *Unclassified.g_Bifidobacterium* (healthy 20x; prediabetic 5x), *B. infantis*, *B. A sp003471165*, *Unclassified.g_Schaedlerella*, *Phocaeicola dorei*, *Unclassified.f_Lachnospiraceae*, *D. formicigenerans*, *Fusicatenibacter saccharivorans*, *B. A sp 900066205*, *A. muciniphila, B. A massiliensis, Unclassified.g_Collinsella*, *B. uniformis*, *F. prausnitzii J, Dialister sp900543165*, *Unclassified.g_Streptococcus, Unclassified.g_Enterococcus B* and *Unclassified.g_Faecalibacterium*, which were all more abundant in healthy fermenta at 0h. Conversely, *F. prausnitzii C, Enterocloster sp000431375, Faecalicoccus acidiformans, S. thermophilus* and *Oliverpabstia intestinalis* were more abundant in prediabetic fermenta and also reduced at 24h. Only the change pattern in *Collinsella sp900764415* differed between fermenta: it was barely detectable at 0h in both groups, and remained so in the prediabetic fermenta, but by 24h had increased by approximately 90 times in the healthy fermenta. Species that became more abundant after colonic fermentation include *Unclassified.C_Gammaproteobacteria* (healthy 236x; prediabetic 1657x); *Unclassified.d_Bacteria* (healthy 90x; prediabetic 83x); and *Enterococcus faecalis* (healthy 90x; prediabetic 1341x). Species that were not detected in either group but appeared differentially post-fermentation include *Paraclostridium bifermentans A* (20 x higher in healthy fermenta); *Unclassified.g_Escherichia* (200x higher in healthy fermenta); *Escherichia coli* (2.5x higher in healthy fermenta); as well as *Unclassified.g_Fusobacterium A*, and *Fusobacterium A ulcerans*. Mean relative abundance at class, family and order levels are available in the Supplementary material (Supplementary material 1, Figure 4 and Supplementary material 2).

**Figure 4.**
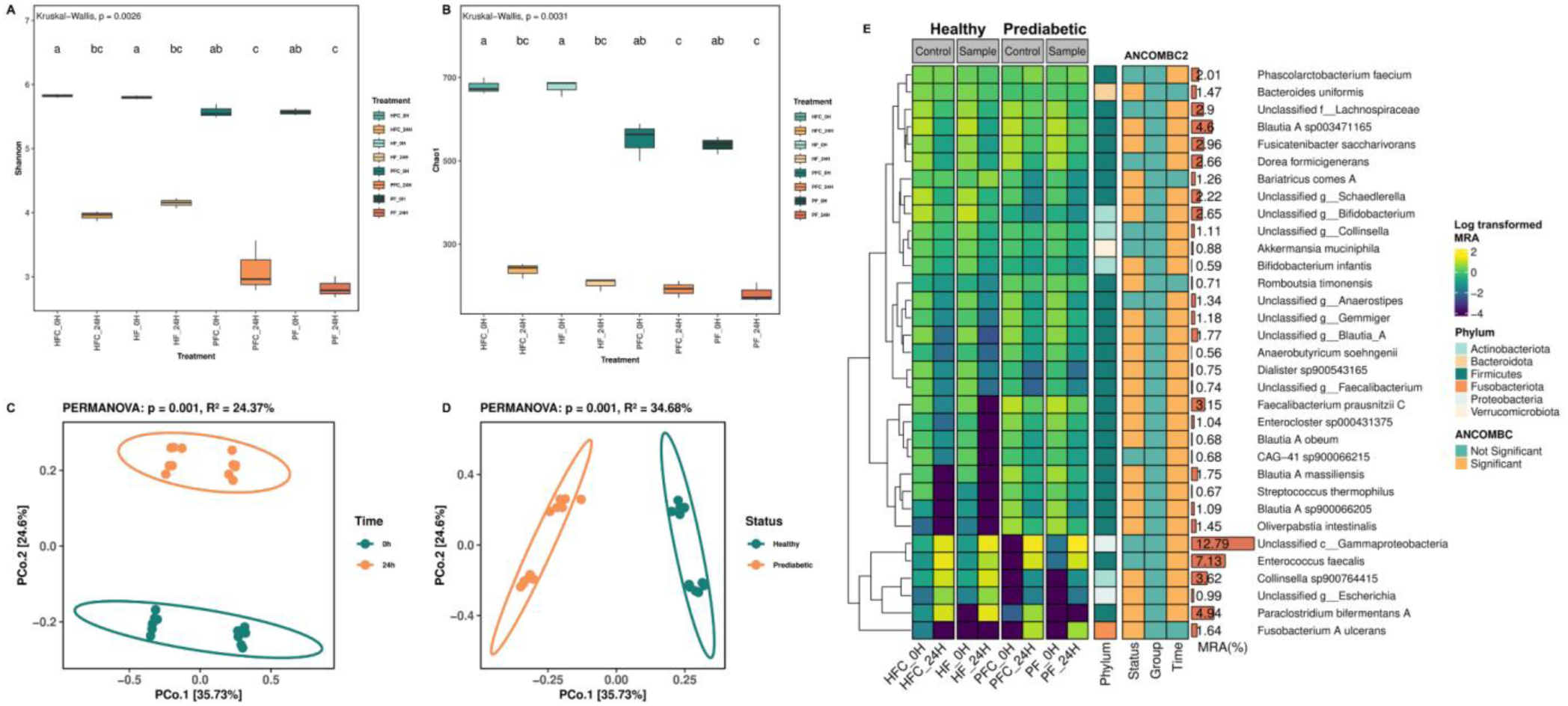
Time and Status affect diversity and abundance of bacterial species in healthy and prediabetic fermenta. Comparison of **α-diversity** represented by (A) Shannon’s (evenness and diversity) and (B) Chao1 (richness) diversity indices; different letters indicate significant differences between samples, as assessed with Dunn’s test. **β-diversity** represented by unweighted UniFrac principal coordinate analysis plots based on ASVs, comparing (C) time points (Time), and (D) healthy and prediabetic fermenta (Status). (E) **Combined clustered/ ANCOM-BC2 heatmap**. From left to right: Clustered heatmap of mean relative abundance (MRA >0.5%) at species and phylum level: Colour of squares in sample columns mapped to abundance; lighter colours indicate higher abundance, while darker colours indicate lower abundance. Abundance of phyla: different colours represent individual phyla. Colour-coding used to visualise significant results of ANCOMBC2 pairwise analysis of bacterial taxa (Status/ Group/ Time). MRA% across all healthy and prediabetic fermenta (*n*=24). Significance level was set at *p*<0.05.

### Quantification of SCFAs during *in vitro* digestion and colonic fermentation

#### Healthy faeces contains significantly higher levels of short-chain fatty acids than prediabetic faeces

Ginger-cayenne tibicos contains high concentrations of SCFAs, especially propionate and acetate. However, *in vitro* digestion of ginger-cayenne tibicos led to significant reductions in all SCFAs: acetate (*p*=0.042); butyrate (*p*=0.0208); propionate (*p*=0.0006); and total SCFA (*p*=0.0098) (Figure 3Ai to iv). As seen in Figure 3Bi to iv, compared to prediabetic faeces, healthy faeces contained significantly higher levels of acetate (*p*<0.0001), butyrate (*p*=0.0104), propionate (*p*=0.0002), and total SCFA (*p*<0.0001).

#### SCFA concentration significantly increased during *in vitro* colonic fermentation

Although colonic fermentation led to an increasing trend across all SCFA concentrations (Figure 3Ci to iv), only some of these changes were significant. Acetate was present in the highest concentrations. Both healthy and prediabetic fermenta samples were not significantly different to their controls at corresponding timepoints, except for acetate concentration in prediabetic fermenta at 8h (*p*=0.0329) (Figure 3Ci). Acetate (Figure 3Ci) significantly increased (*p*=0.0071) in healthy fermenta between 0h and 12h, whilst prediabetic fermenta showed significant increases between 0 and 8h (*p*=0.110), and 4 and 8h (*p*=0.0043). Butyrate (Figure 3Cii) increased significantly between 4 and 24h (*p*=0.0189), and 8 and 24h (*p*=0.0187) in the healthy fermenta, with no significant shifts in the prediabetic fermenta. Propionate (Figure 3Ciii) levels in the healthy fermenta increased significantly between various timepoints, but most importantly was significantly higher at 24h (*p*=0.0499) compared to the start of fermentation. In the prediabetic fermenta, propionate was significantly increased between 0 and 8h (*p*=0.0184), and 4 and 8h (*p*=0.0227). Total SCFAs in healthy fermenta were significantly increased between 0 and 12h (*p*=0.0041), whilst in prediabetic fermenta, concentrations were significantly higher at 8h vs. 0h (*p*=0.0082) and 4 h (*p*=0.0107). When comparing the two groups, propionate was significantly higher in healthy fermenta at 12 h (*p*=0.0004) (Figure 3Ciii). Overall, repeated measures ANOVA showed that fermentation time had a significant impact on SCFA concentrations.

### Diversity, differential and correlation analyses

#### β-diversity of healthy and prediabetic faeces differs and is affected by *in vitro* colonic fermentation

α– and β-diversity of ginger-cayenne tibicos did not differ significantly before and after *in vitro* digestion (Supplementary material 1, Figure 5A, B). There was no significant difference in alpha or beta diversity between healthy and prediabetic faecal samples, likely due to the small sample size (*n*=3 donors per group) (Supplementary material 1, Figure 5C, D).

**Figure 5.**
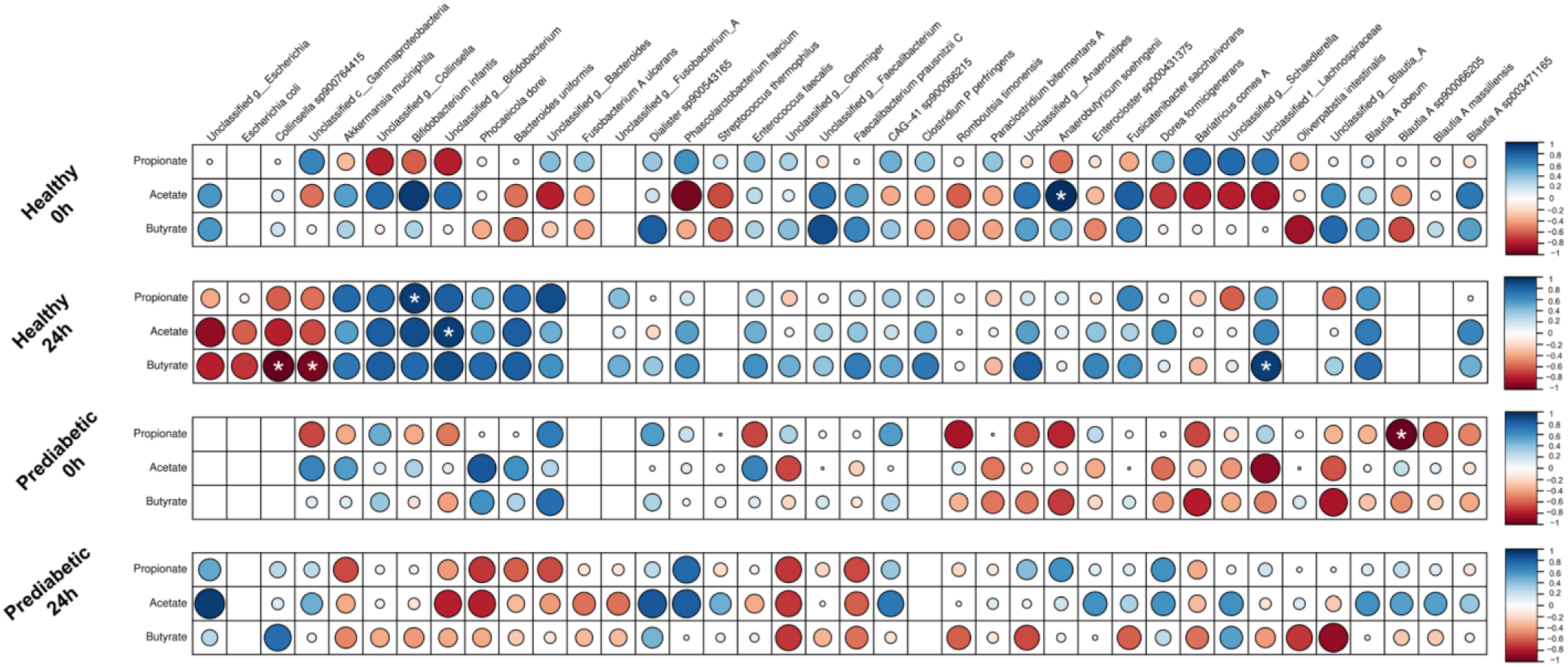
Spearman’s correlation heatmap between bacterial species (MRA >0.5%) and faecal SCFA concentrations in healthy and prediabetic fermenta at 0h and 24h of colonic fermentation: Blue dots represent positive *r* values/ correlation; red dots represent negative *r* values/ correlation; empty squares indicate absent species; colour intensity and size of dots indicate the magnitude of correlation; and * indicates significantly correlated pairs (FDR-adjusted *p* value <0.1).

Both the healthy and prediabetic fermenta differed in α-diversity between 0h and 24h of *in vitro* colonic fermentation, as observed by Shannon’s (*p*=0.0026) and Chao1 (*p*=0.0031) diversity indices (Figures 4A, B respectively). However, there were no significant differences between the healthy vs. prediabetic fermenta, or the controls vs. samples. β-diversity was assessed using unweighted UniFrac in principal coordinate analysis (PCoA) plots, demonstrating significant differences only in Time (0h vs 24h) and Status (healthy vs. prediabetic fermenta) (both PERMANOVA, *p*=0.001) (Figures 4C, D respectively).

#### Time and Status affect differential shifts in bacterial species

A clustered heatmap (Figure 4E) was used to further understand differential shifts in relative abundance of phyla and species. Across all samples, the dominant phylum by far was *Firmicutes*, followed by *Actinobacteriota* and *Proteobacteria*, with a low abundance of *Bacteroidota* and *Fusobacteriota*. On a species level, *Proteobacteria Unclassified.c_Gammaproteobacteria* (12.79%) dominated, followed by *Firmicutes E. faecalis* (7.13%), *P. bifermentans* (4.94%) and *B. A sp003471165* (4.6%). The healthy and prediabetic fermenta were affected differently by 24h of colonic fermentation. A drop in *Firmicutes* in both groups was observed, but there was a greater reduction in the healthy fermenta. An increase in the abundance of *Unclassified.c_Gammaproteobacteria* was also observed but was greater in the prediabetic fermenta. Compared to the prediabetic fermenta, healthy fermenta saw larger increases in *E. faecalis, Unclassified.g_Collinsella, Unclassified.g_Escherichia* and *P. bifermentans A* abundance, with larger reductions in many *Firmicutes*, including *F. prausnitzii C, E. sp000431375, S. thermophilus, CAG-41 sp900066215,* and several *Blautia A* species.

ANCOM-BC2 was used to further analyse the precise differences in microbial species between the two groups (Figure 4E). The results confirmed our earlier findings, in that none of the involved species were significantly different between the controls and samples (Group), but the majority were significantly different in relation to Time and Status. Specifically, there were no significant differences (*p*<0.05) between the groups in *P. faecium, A. muciniphila, D. formicigenerans, Unclassified.f_Lachnospiraceae, Unclassified.g_Collinsella,, Unclassified.g_Anaerostipes, CAG-41 sp900066215, O. intestinalis* and the dominant species *Unclassified.c_Gammaproteobacteria* and *E. faecalis.* In relation to changes before and after colonic fermentation, all observed species shifted significantly, except for *B. uniformis, Bariatricus comes A, Romboutsia timonensis* and *F. A ulcerans*.

#### Correlation patterns of SCFAs and faecal microbiota differed between groups and were affected by colonic fermentation

Given the known relationship between prediabetes and obesity on the gut microbiota and SCFA production, we analysed the associations between faecal SCFAs and relatively abundant bacterial species (Figure 5). The healthy fermenta had different correlation patterns and more overall significant correlations compared to the prediabetic fermenta, and colonic fermentation shifted correlation patterns in both groups. The healthy fermenta at 0h only displayed a significant positive correlation between *Anaerobutyricum soehngenii/*acetate (*q*=0). At 24h, correlations were generally strengthened in either a positive or negative direction. The correlation between *B. infantis/* propionate went from being strongly negative, to significantly positive (*q*=0.09), whilst this species was also (not significantly) positively correlated with acetate and butyrate. *Unclassified.g_Bifidobacterium* went from being strongly negatively correlated with propionate, to positive correlations with all SCFAs, significantly so with acetate (*q*=0.09). *Unclassified.f_Lachnospiraceae* had a strong negative correlation with acetate, but by 24h was strongly positively correlated with all SCFAs, although only butyrate was significant (*q*=0.09). *Colinsella sp900764415* and *Unclassified.c_Gammaproteobacteria* both had relatively weak correlation patterns at 0h, but by 24h had strong negative correlations with all SCFAs, significantly so with butyrate (*q*=0 and *q*=0.09 respectively). Prediabetic fermenta had only one significant negative correlation (*q*=0.03) at 0h, *B. A sp900066205*/propionate, and none at 24h, although colonic fermentation did change overall correlation patterns.

## Discussion

In *in vitro* digestion/ colonic fermentation studies using healthy faeces, tibicos has been shown to contain probiotic levels of microorganisms that survive GI transit, beneficially shift faecal microbiota composition, and increase microbial SCFA production ^(8,17,25)^. No such studies comparing tibicos’ effects on healthy vs. prediabetic faecal microbiota have been conducted. Our study showed that healthy and prediabetic faecal microbiota have distinct microbial compositions and functional capabilities and are affected by gastrointestinal transit and ginger-cayenne tibicos in different ways.

### Ginger-cayenne tibicos contains potentially beneficial bacteria and SCFAs that are reduced by *in vitro* digestion

The microbial composition of the tibicos was similar to that of other studies ^(13,42–44)^, with LAB species such as *Liquorilactobacillus hordei*, *Liquorilactobacillus nagelii,* and *Leuconostoc mesenteroides* dominating. As expected, due to the low pH environment and enzymatic activity involved, *in vitro* digestion significantly reduced (*p*=0.001) the bacterial content of ginger-cayenne tibicos ^(8)^. Slight upwards and downwards changes in mean relative abundance of LAB in tibicos were observed, confirming the ability of some species to withstand GI transit ^(19)^. Some potentially beneficial species only appeared post-digestion, such as *A. muciniphila* and *F. prausnitzii*. These shifts are likely due to the enzymatic release of prebiotics and antioxidant phenolic compounds from ginger and cayenne ^(8,45–47)^. As usually observed ^(17)^, the tibicos in our study initially had high acetate, propionate and butyrate levels. Due to the reduction in bacterial content and shifts in relative abundance of butyrate producers, these SCFA levels were all significantly reduced (*p*<0.02) post-digestion. Overall, *in vitro* digestion affected the tibicos microbial community by reducing bacterial numbers, shifting relative abundance, thus lowering SCFA concentrations.

### Healthy and prediabetic faeces have distinct microbial composition and function which are differentially affected by *in vitro* colonic fermentation

Compared to healthy participants, faeces from prediabetic and obese participants are known to have reduced microbial diversity and differing abundances of bacterial genera involved in metabolic and inflammatory processes, with subsequent functional consequences ^(48–50)^. However, Letchumanan et al. ^(49)^ found, in their systematic review of observational studies that, despite a general pattern of reduced diversity, differences in microbial composition were inconsistent across studies. Overall, our findings concurred with the current evidence.

In our study, prediabetic faeces was more abundant in certain genera/species, and less abundant in others. Most of our findings concurred with other studies as per Letchumanan et al.’s systematic review ^(49)^. Notably, we found that prediabetic faeces was 15 times less abundant in *Bifidobacterium*, which is a common finding in prediabetic and T2DM participants ^(50,51)^. In animal models, *Bifidobacterium* species have been observed to positively shift faecal microbiota composition, improve glycaemic and blood lipid control and reduce inflammation ^(52–54)^. *Prevotella* was 100 times less abundant in prediabetic faeces, as observed in Egshatyan et al. ^(55)^ and Zhang et al. ^(56)^. Associated with a plant/ fibre-rich dietary pattern, this important intestinal commensal has high genetic diversity ^(57)^, with different strains having beneficial effects on glucose metabolism ^(58,59)^, whilst others promote development of MetS and obesity ^(60)^. The GTDB database we used only identified *P. sp900551275* in our samples, which is known to be more prevalent in nonindustralised populations with fibre-rich dietary patterns ^(61)^. We also found that *P. faecium*– a potential biomarker for the early detection of prediabetes and T2DM– was less abundant in prediabetic faecal samples, which concurs with the current literature ^(62–64)^. *Bacteroides* was more abundant in prediabetic faeces, which is expected, as *Bacteroides* is associated with a higher risk of developing T2DM ^(65)^. However, *B. uniformis* was more abundant in healthy faeces, as also observed in Allin et al. ^(66)^. Diabetic animal models indicate that *B. uniformis* improves glucose tolerance, insulin secretion, postprandial glycaemic control ^(67)^, and lipid metabolism ^(68)^. As an example of the diversity of this genus, *Bacteroides fragilis*, which we did not observe in our study, is more abundant in prediabetic faecal samples ^(69)^. This highlights the importance of conducting research at a species/ strain level. *Gemmiger*/ *Unclassified.g_Gemmiger*– butyrate producers involved in inflammatory responses– were more abundant in the prediabetic group, which seems counterintuitive as the presence of this genus/ species is negatively correlated with HbA1c and fasting blood glucose ^(70)^. However, Zeng et al. ^(71)^ found that obese participants with different metabolic abnormalities shared gut microbiota biomarkers of disease, including *Blautia, Dorea* and *Gemmiger*. As such, our finding of less abundance of *Dorea/ D. formicigenerans* in the prediabetic group is surprising: this genus/species is typically more abundant in individuals with prediabetes ^(66,72,73)^, and has a positive correlation with impaired fasting glucose ^(74)^. Unusually, we found that *A. muciniphila* was four times more abundant in prediabetic faeces, in contrast to other studies ^(56,66,75)^. In animal and human studies, *A. muciniphila* is negatively correlated with HbA1c in T2DM ^(76)^ and obesity ^(77,78)^, and has been shown to improve insulin sensitivity, reduce fasting blood glucose levels, and reduce cholesterol levels ^(78–81)^. We also found that *F. prausnitzii C* was five times more abundant, and *F. prausnitzii J* five times less abundant, in prediabetic faeces. This finding concurs with Allin et al. ^(66)^ in that two different strains of *F. prausnitzii* were found to be more or less abundant in prediabetic faeces. However, *F. prausnitzii* was less abundant in prediabetic faeces in other studies ^(69,75,82)^. This important commensal species, usually highly abundant in healthy adults, is a major butyrate producer in the gut ^(83)^, and has been shown to decrease intestinal permeability, as well as produce antiinflammatory molecules ^(84–86)^. Prediabetic and diabetic mice studies indicate that *F. prausnitzii* reduces fasting blood glucose, lowers HbA1c and improves glucose response ^(87)^. As such, it has generated interest as a potential gut microbiota-targeted treatment for noncommunicable cardiometabolic diseases ^(85)^. These departures from the literature may be due to the dietary pattern of our participants leading up to the faecal collection date, as short-term dietary changes can rapidly alter gut microbiota ^(2,88)^, as well as interindividual variation in baseline gut microbiota composition. Our findings must be considered in the context of our small sample size.

Our study found that fermentation Time and Status (i.e. healthy vs. prediabetic) had the biggest impact on microbial community abundance and diversity. Ginger-cayenne tibicos was able to modify both prediabetic and healthy faecal microbial communities, although the effects were small. The similar decline and increase in particular bacterial taxa in both fermenta during *in vitro* colonic fermentation were in line with Goya-Jorge et al. 2022’s study of human adult gut microbiota in a static colon model. Though following similar patterns of change, the baseline faecal microbiota of each group differentially affected fermentation of tibicos’ dietary substrates, and the interactions between tibicos-associated microbes and commensals ^(89)^. This led to factorial differences in the upwards or downwards trends in relative abundance (Figures 2A to C; Figure 4E). For instance, both fermenta, *in vitro* colonic fermentation with ginger-cayenne tibicos led to reductions in *Firmicutes*, *Bacteroidetes*, and *Verrucomicrobiota*, with large increases in *Proteobacteria*. This is a common finding in such studies, with specific variations depending on the substrates and populations involved ^(17,89,90)^. Compared to the healthy fermenta, prediabetic fermenta had a larger reduction in *Firmicutes* abundance, with the converse being true for *Bacteroidetes*, and a much larger increase in *Proteobacteria.* Prediabetic fermenta had smaller reductions in the abundance of beneficial genera/ species, including *Blautia A, Bifidobacterium/ Unclassified.g_Bifidobacterium, Dorea,* and *Fusicantenibacter*. These findings differ from Calatayud et al.’s ^(17)^ study which observed enrichment of *Bifidobacterium* in healthy faeces with the application of pasteurised kefir. Healthy fermenta saw large increases in pathogenic *Paraclostridium* and *Escherichia,* whilst these both remained at 0% in prediabetic fermenta. *Collinsella/ C. sp900764415* positively correlate with higher circulating insulin and cholesterol and are increasingly abundant with low-fibre diets ^(91)^. In our study, this genera/species increased by 90x in healthy fermenta, while it remained undetectable/ was reduced in prediabetic fermenta. A similar trend was seen with *P. bifermentans A* and *Unclassified.g_Escherichia/ E. coli*. These shifts could indicate that tibicos exerts more protective effects in prediabetic fermenta. However, in contrast, prediabetic fermenta displayed larger increases in *Unclassified.c_Gammaproteobacteria*, *Unclassified.d_Bacteria* and *Enterococcus/ E. faecalis,* all of which comprise many pathogenic species. Beta diversity was significantly (PERMANOVA *p*=0.001) different between the fermenta and became more dissimilar over fermentation time (PERMANOVA *p*=0.001). α-diversity was only significantly affected by colonic fermentation time (Shannon’s, Kruskal-Wallis test, *p*=0.0026; Chao1, Kruskal-Wallis test, *p*=0.0031), although generally the healthy fermenta had higher biodiversity than the prediabetic fermenta. These findings concur with Chang et al.’s ^(50)^ study comparing the gut microbiota composition of prediabetic and healthy participants. Calatayud et al.’s ^(17)^ *in vitro* digestion study of water kefir also observed that the main effects on the diversity of microbial communities were fermentation time and donor, and not kefir treatment, compared to control. We also found no differences to control for all diversity indices, suggesting that although ginger-cayenne tibicos ‘consumption’ leads to shifts in microbial abundance, the dosage may not be sufficient to engender significant shifts in microbial diversity. Ginger-cayenne tibicos differentially affects faecal microbiota of healthy and prediabetic participants, and may have more beneficial effects on prediabetic faeces, but studies with larger sample sizes and tibicos dosages are needed.

Colonic microbial fermentation of dietary fibre and phenolic compounds results in overall increases in the major SCFAs acetate, butyrate and propionate in faeces. These beneficial metabolites, especially butyrate, act as primary energy sources for colonocytes ^(92)^. SCFAs strengthen intestinal barrier integrity, regulate immune function and inflammatory responses, with impacts on glucose metabolism, appetite regulation and lipid metabolism ^(92–94)^. Firstly, we found that at baseline, healthy faeces contained significantly (*p*<0.0001) higher levels of acetate, butyrate and propionate, compared to prediabetic faeces. Other studies have shown that faecal SCFA concentrations, particularly butyrate and propionate, are lower in prediabetic individuals compared to healthy controls ^(72,92)^, which may contribute to insulin resistance and impaired glucose control ^(94,95)^. This is likely due to the reduced diversity of prediabetic faecal microbiota, and decreased abundance of butyrate-producing bacteria ^(92)^. *In vitro* colonic fermentation with ginger-cayenne tibicos led to overall significant (*p*<0.05) increases in all SCFAs over time in both healthy and prediabetic faecal microbiota, although the patterns of change differed. Acetate, which is produced by many of the dominant bacterial phyla during colonic fermentation, is found to be present in the highest concentrations in faeces ^(96)^ and is commonly metabolised by other gut bacteria to produce butyrate, with beneficial effects on the gut and beyond ^(97)^. We found that acetate was present at the highest concentrations, compared to propionate and butyrate. This was similar to findings in studies exploring the *in vitro* and *in vivo* effects of water kefir ^(17)^, lactofermented vegetables ^(98)^, and milk kefir ^(99)^, on healthy faecal microbiota. The healthy fermenta had more significant increases in concentration by 24h, especially in butyrate and propionate. However, the only significant difference between the groups was that healthy fermenta contained higher levels of propionate (*p*=0.0004) at 12h. With higher baseline SCFAs, and larger increases in SCFAs post-fermentation, it was not unexpected to find that healthy fermenta had more significant correlations between bacterial species and SCFAs than the prediabetic fermenta. After 24h, we observed significant positive correlations between *Unclassified.g_Bifidobacterium/* acetate. *Bifidobacterium* species convert dietary carbohydrates into acetate, which can be converted to butyrate by other gut commensals, modulating gut microbiota composition and intestinal homeostasis, thus exerting anti-hyperglycaemic, antiinflammatory and antipathogenic effects on the host ^(100–102)^. We also observed significant positive correlations between *Unclassified.f_Lachnospiraceae*/ *Colinsella sp900764415/ Unclassified.c_Gammaproteobacteria* butyrate post fermentation, indicating an increase in butyrate producers, with beneficial downstream effects on glucose metabolism. Calatayud et al. ^(17)^ demonstrated that pasteurised water kefir administered to healthy faeces increased the *in vitro* production of SCFAs and anti-inflammatory cytokines IL-10 and IL-1β, as well as activating key signalling molecule NF-κB, all of which promote intestinal barrier homeostasis. This indicates that, in our study, healthy fermenta contained more SCFA-producing bacteria that were able to metabolise ginger-cayenne tibicos components, resulting in higher SCFA concentrations.

Our study contributes to the literature on the effects of tibicos on faecal microbiota and its potential as a complementary management tool in prediabetes. As far as we can surmise, the only other *in vitro* digestion study using human faecal microbiota with water kefir is Calatayud et al. ^(17)^. However, our study is the first to compare prediabetic and healthy participants in this context, moving one step closer to establishing tibicos’ potential role in the management of prediabetes. That said, a major limitation of our study was the small sample size, due to the difficulty of recruiting eligible prediabetic/ obese participants. One reason for this was our exclusion of participants using antidiabetic medications such as metformin and GLP-1 agonists, as these agents are known to influence gut microbiota composition and function ^(103)^, and thus obscure potential effects of other interventions. Most prediabetic and obese patients are prescribed such medications upon diagnosis, and as such, it was difficult to find medication-naive participants. Interindividual variability in gut microbiota, dietary patterns leading up to the study, as well as the ethnicity of our participants may also have had an impact on our findings. Personalisation was beyond the scope of our study but is an ideal to work towards in future diet-microbiome research. Potential measurement errors in some samples, likely due to sample storage conditions, were identified and still included. Due to the cost of shotgun metagenomics, we opted for long-read 16S rRNA sequencing for (sub)species level differentiation ^(104)^. As such, although our study indicates a rationale for more studies of tibicos for use in prediabetes management, the translatability of our findings is limited. For future studies, we suggest larger sample sizes, dosage considerations, dietary controls and more ethnically diverse participants, as well as personalisation.

## Conclusions

Prediabetic and healthy faecal microbiota differ in composition and function, with prediabetic faeces being less abundant in beneficial butyrate producers and SCFAs. Ginger-cayenne tibicos, containing live microorganisms in probiotic numbers, dietary fibres, and phenolic compounds, differentially affects prediabetic and healthy faecal microbiota during *in vitro* colonic transit. Fermentation time and status have the largest impact on microbial community abundance and diversity. Tibicos may have more beneficial effects on prediabetic faeces composition, with less reduction of beneficial bacterial species. Dosage studies and larger sample sizes are required to ascertain clinical validity.

## Supplementary material

Supplementary material is provided here: Supplementary material 1 and 2.

## Acknowledgements

The authors would like to thank Professor Elif Ekinci and Nupoor Tomar at the Centre for Education and Research in Diabetes and Obesity, Austin Health for coordinating collection of faeces; and Siyao Liu and Dillani Putri Ramadhaningtyas for assistance in HPLC.

## Authorship

MC and KH conceived and designed the study. LN and JW conducted the laboratory experiments and data analysis with MC. XR analysed the microbiome data with MC. MC wrote and revised the manuscript with editorial input from KH. All authors approved the manuscript.

## Data availability statement

Raw sequencing reads have been deposited at the National Centre for Biotechnology Information Sequence Read Archive here: https://www.ncbi.nlm.nih.gov/bioproject/PRJNA1242392. All other data presented in the study are included in the article and supplementary material.

