## Supplemental material 1 for "Tibicos differentially affects faecal microbiota composition and short-chain fatty acid production in prediabetic and healthy adults in a simulated digestive tract"

**Supplementary material 1**

**Bacterial content of tibicos, faecal samples and fermenta**

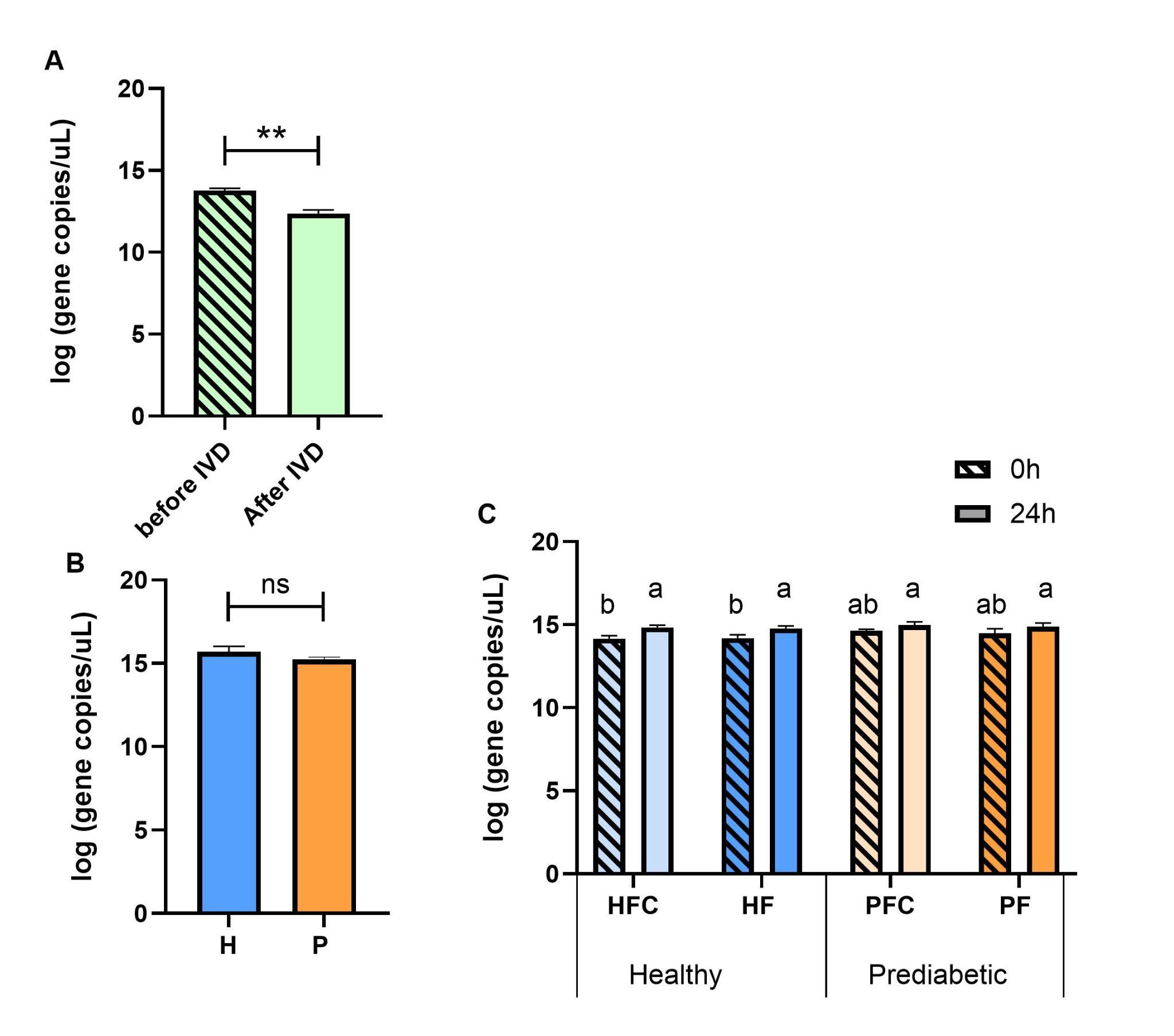

**Figure S1.** Total number of bacterial gene copies in **(A)** tibicos samples before and after *in vitro* digestion (IVD); **(B)** healthy (H) and prediabetic (P) faecal samples; **(C)** *in vitro* fermentation samples of healthy (HF) and prediabetic (PF) group and their fermentation controls at 0 and 24 h of fermentation. HFC and PFC controls contained the *in vitro* digestion control samples with MilliQ water instead of tibicos. Results are represented as mean log-transformed gene copies per µL of sample. ** Statistically significant difference (*p*≤ 0.05); ^a-b^ Different superscript lowercase letters for *in vitro* fermentation samples indicate significant differences in the mean log GCN.

**Class, order and family of tibicos, faecal samples and fermenta**

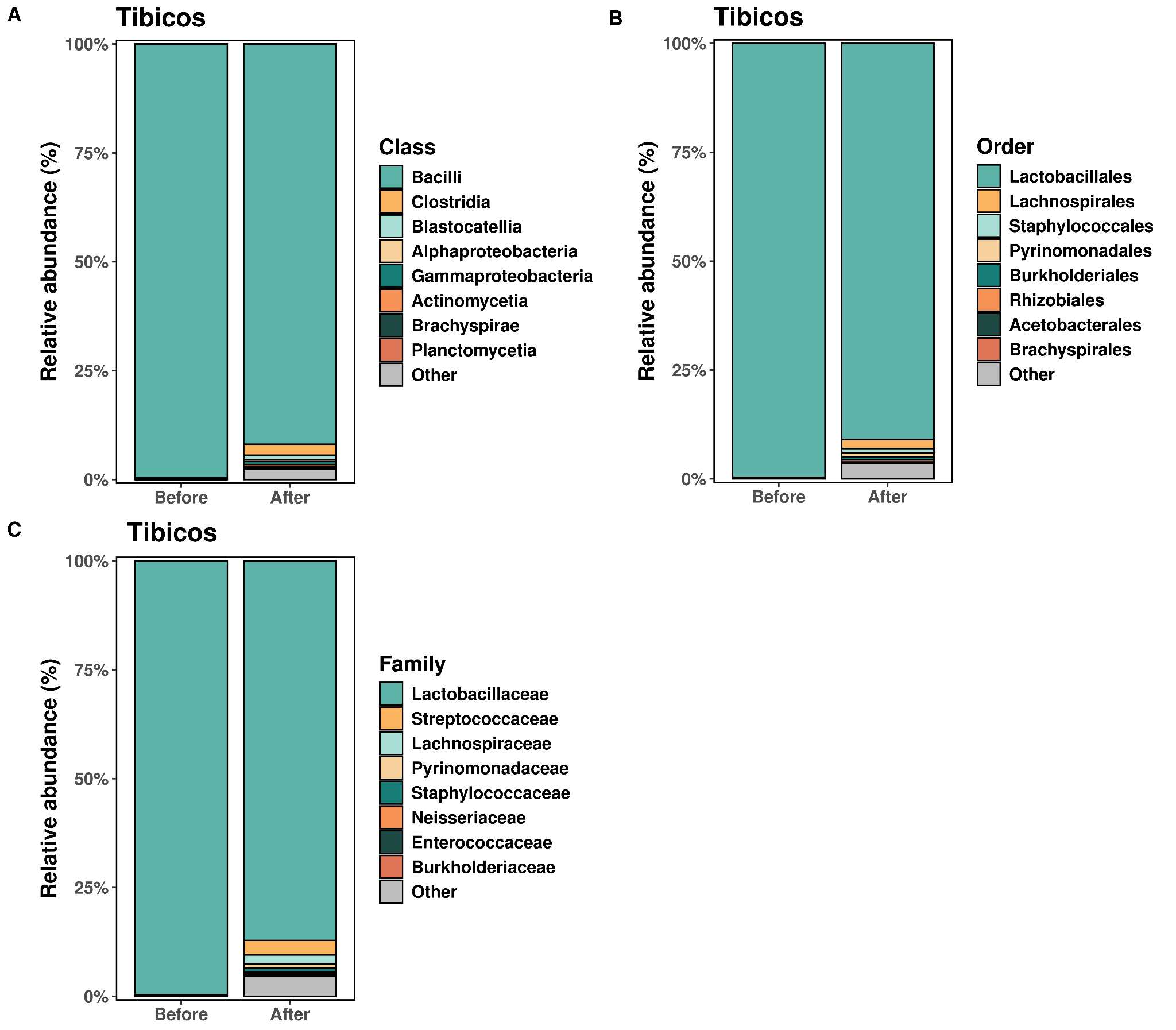

**Figure S2.** The microbial composition of ginger-cayenne tibicos before and after *in vitro* digestion. Microbial taxa in ginger-cayenne tibicos are classified by **(A)** class, **(B)** order, and **(C)** family levels, and ordered according to abundance. Results are expressed as mean ± SD (*n*=3).

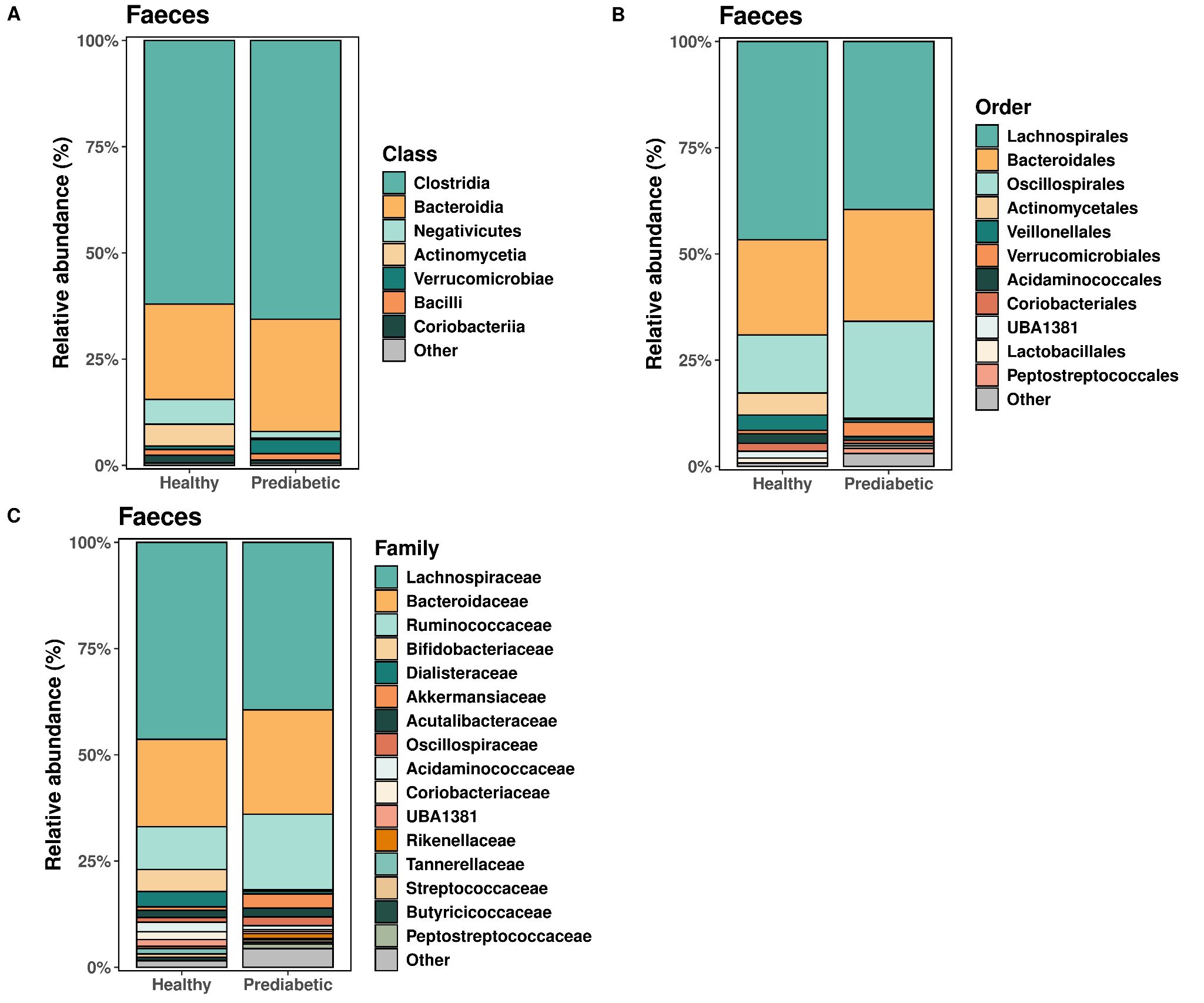

**Figure S3.** Microbial composition of healthy and prediabetic faecal microbiota Microbial taxa in healthy and prediabetic faecal samples are classified by **(A)** class, **(B)** order, and **(C)** family levels, and ordered according to abundance. Results are expressed as mean ± SD (*n*=3).

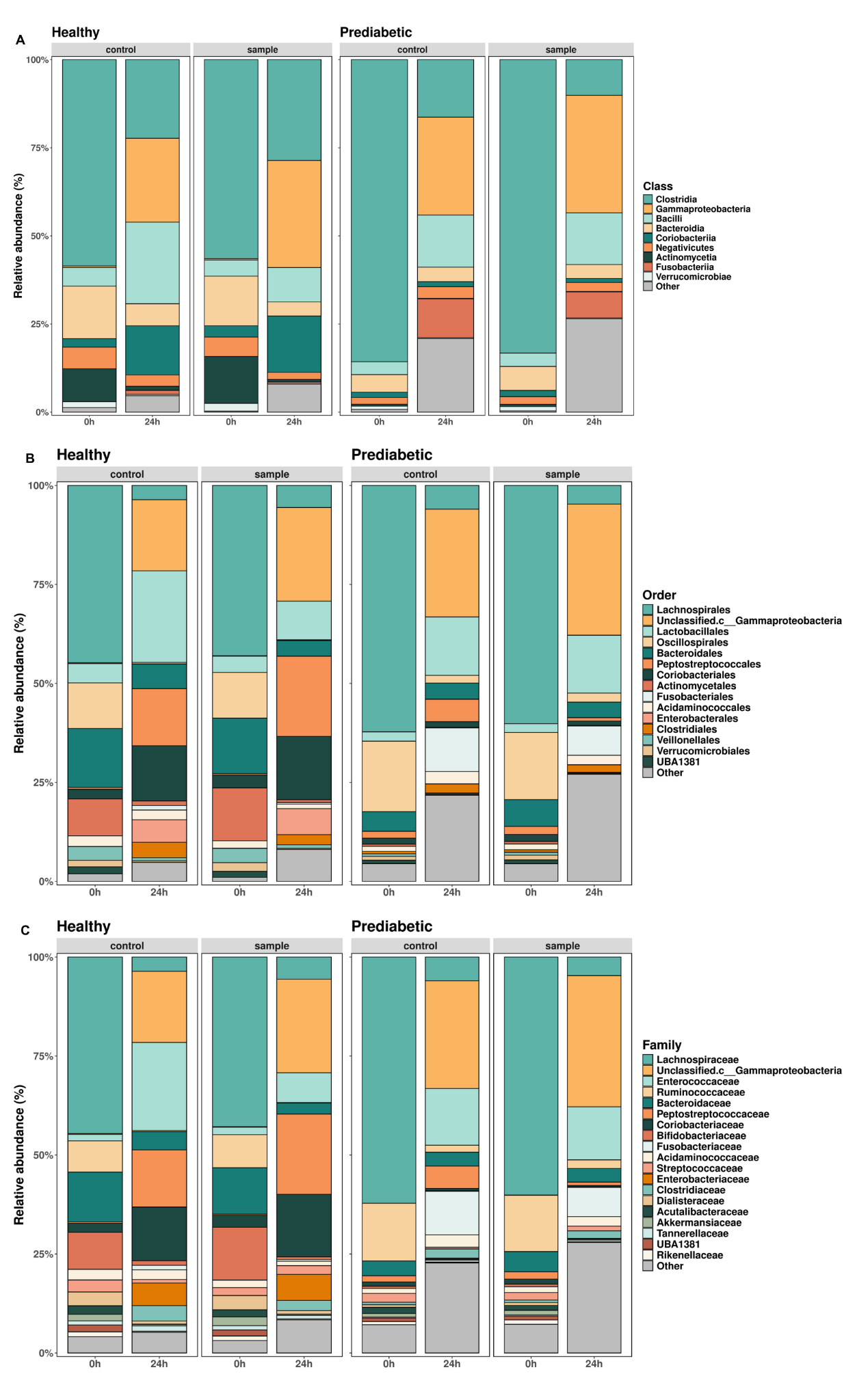

**Figure S4.** Comparison of changes in healthy and prediabetic faecal microbiota composition during *in vitro* colonic fermentation. Relative abundance plots were generated using the most abundant bacterial taxa. Microbial taxa are classified by (A) class, (B) family, and (C) order levels. Healthy and prediabetic samples: fermentation media + faecal slurry + tibicos digesta from IVD; healthy and prediabetic controls: fermentation media, faecal slurry and control digesta from IVD. Results expressed in mean ± SD (*n*=3).

**Diversity indices of tibicos and faecal samples**
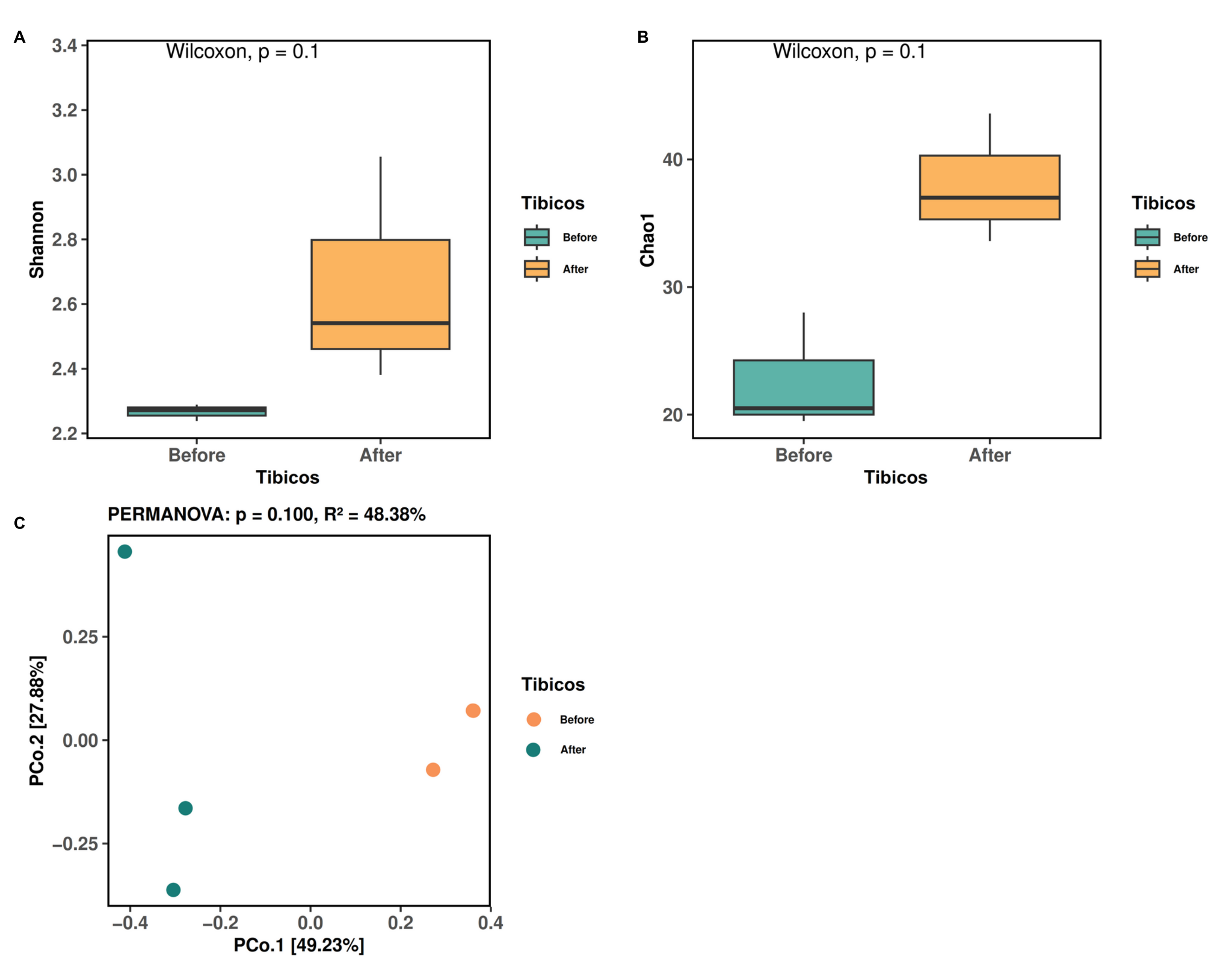

**Figure S5.** α- and β- diversity of ginger-cayenne tibicos before and after *in vitro* digestion. Comparison of **α-diversity** represented by **(A)** Shannon’s (evenness and diversity) and **(B)** Chao1 (richness) diversity indices; differences assessed using Wilcoxon’s signed-rank test. **(C)** **Β-diversity** represented by unweighted UniFrac principal coordinate analysis plots based on ASVs, before and after *in vitro* digestion. Significance level was set at *p*<0.05.

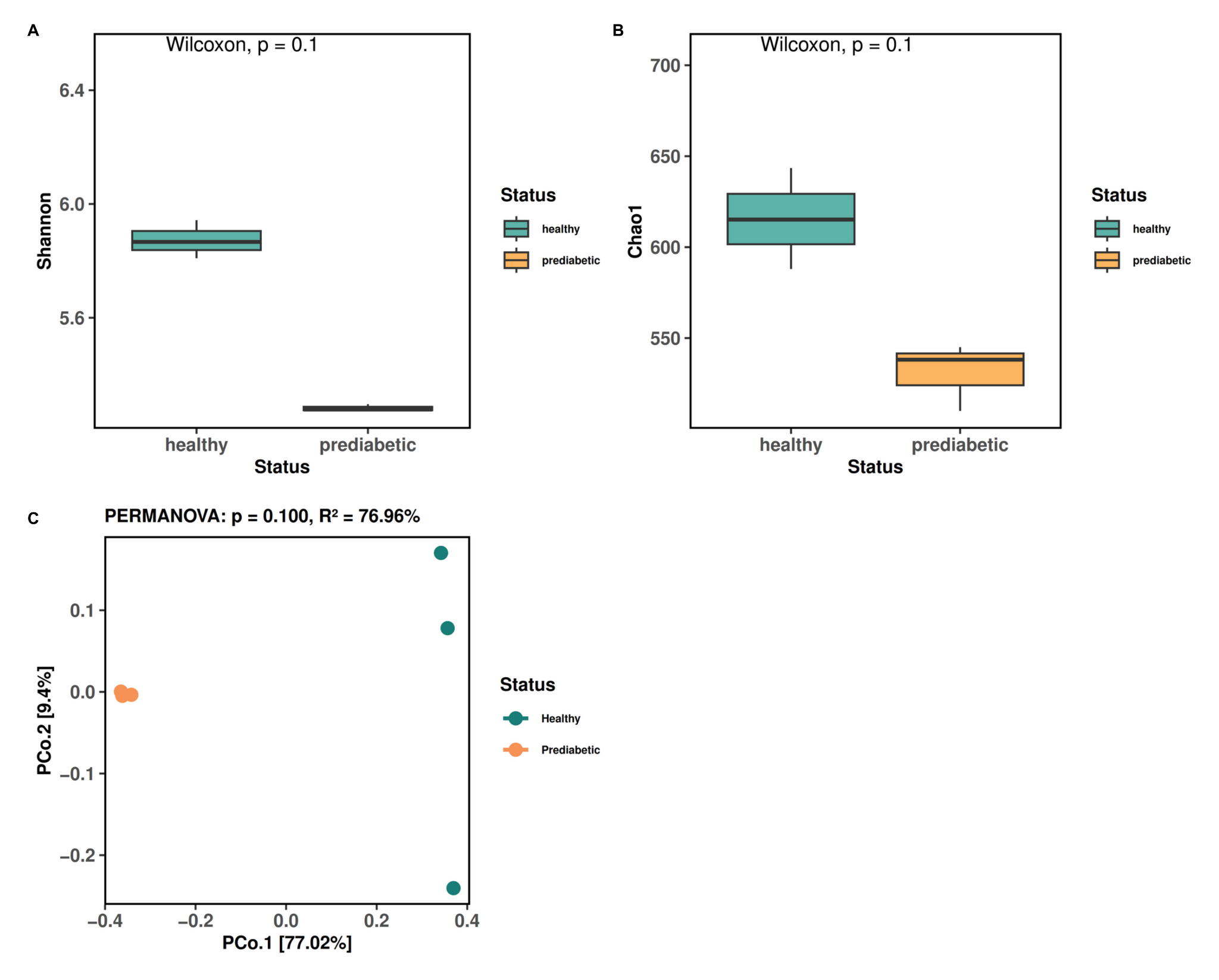

**Figure S6.** α- and β- diversity of healthy and prediabetic faecal samples. Comparison of **α-diversity** represented by **(A)** Shannon’s (evenness and diversity) and **(B)** Chao1 (richness) diversity indices; differences assessed using Wilcoxon’s signed-rank test. **(C)** **Β-diversity** represented by unweighted UniFrac principal coordinate analysis plots based on ASVs, of healthy and prediabetic faecal samples. Significance level was set at *p*<0.05.

***Firmicutes/ Bacteroidetes* ratio before and after *in vitro* colonic fermentation**

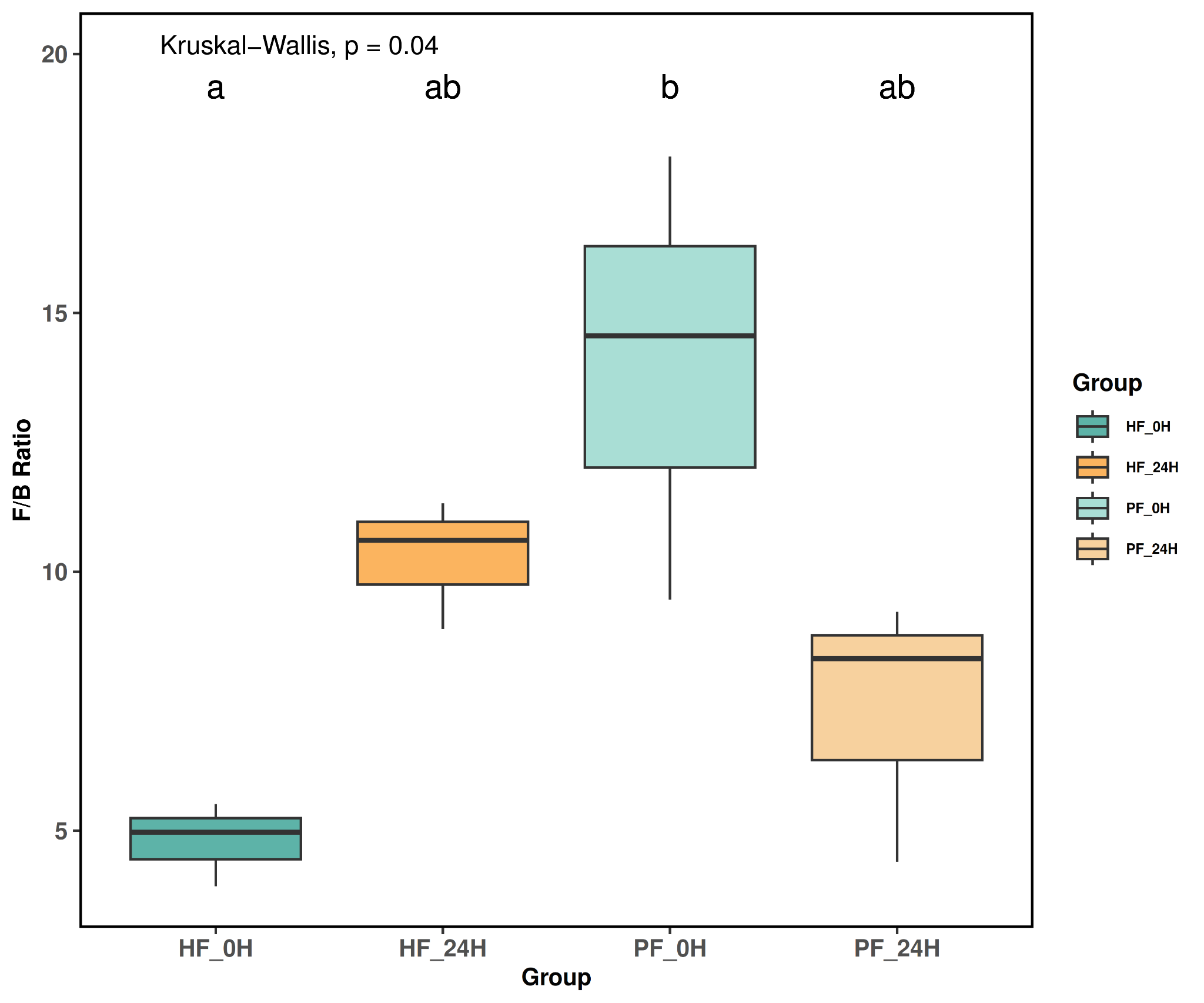

**Figure S7.** *Firmicutes/ Bacteroidetes* (F/B) plot of healthy and prediabetic fermenta before and after *in vitro* colonic fermentation. Different letters indicate significant differences between samples, as assessed with the Kruskal-Wallis test. Significance level set at *p*<0.05.
